# The 3’-untranslated regions of yeast ribosomal protein mRNAs determine paralog incorporation into ribosomes and recruit factors necessary for specialized functions

**DOI:** 10.1101/2024.03.18.585503

**Authors:** Raman K. Singh, Robert A. Crawford, Dheerendra P. Mall, Graham D. Pavitt, Jeffrey E. Gerst

## Abstract

Ribosome heterogeneity arises via the differential incorporation of ribosomal protein (RP) paralogs, post-transcriptionally modified rRNA, post-translationally modified RPs, and ribosome-associated proteins (RAPs) into ribosomes. This has led to the hypothesis that heterogeneous or “specialized” ribosomes, which translate specific mRNA subsets, confer key roles in cell growth and development. While proven examples of functional ribosome heterogeneity in eukaryotes exist, there is no comprehensive analysis of specialized ribosome formation. We employed yeast RP paralog deletion libraries and high-throughput screening to investigate the functional specificity and redundancy between paralogs under various growth conditions. Composition and translatome analyses verified paralog specificity in the assembly and function of ribosomes specialized for growth on different carbon sources, and identified a novel RAP required for the efficient translation of peroxisomal proteins. Importantly, we also show that the mechanism by which specific RP paralogs incorporate into ribosomes requires their unique 3’-untranslated regions to yield ribosomes that differ in composition and function.

**Graphical Abstract:** 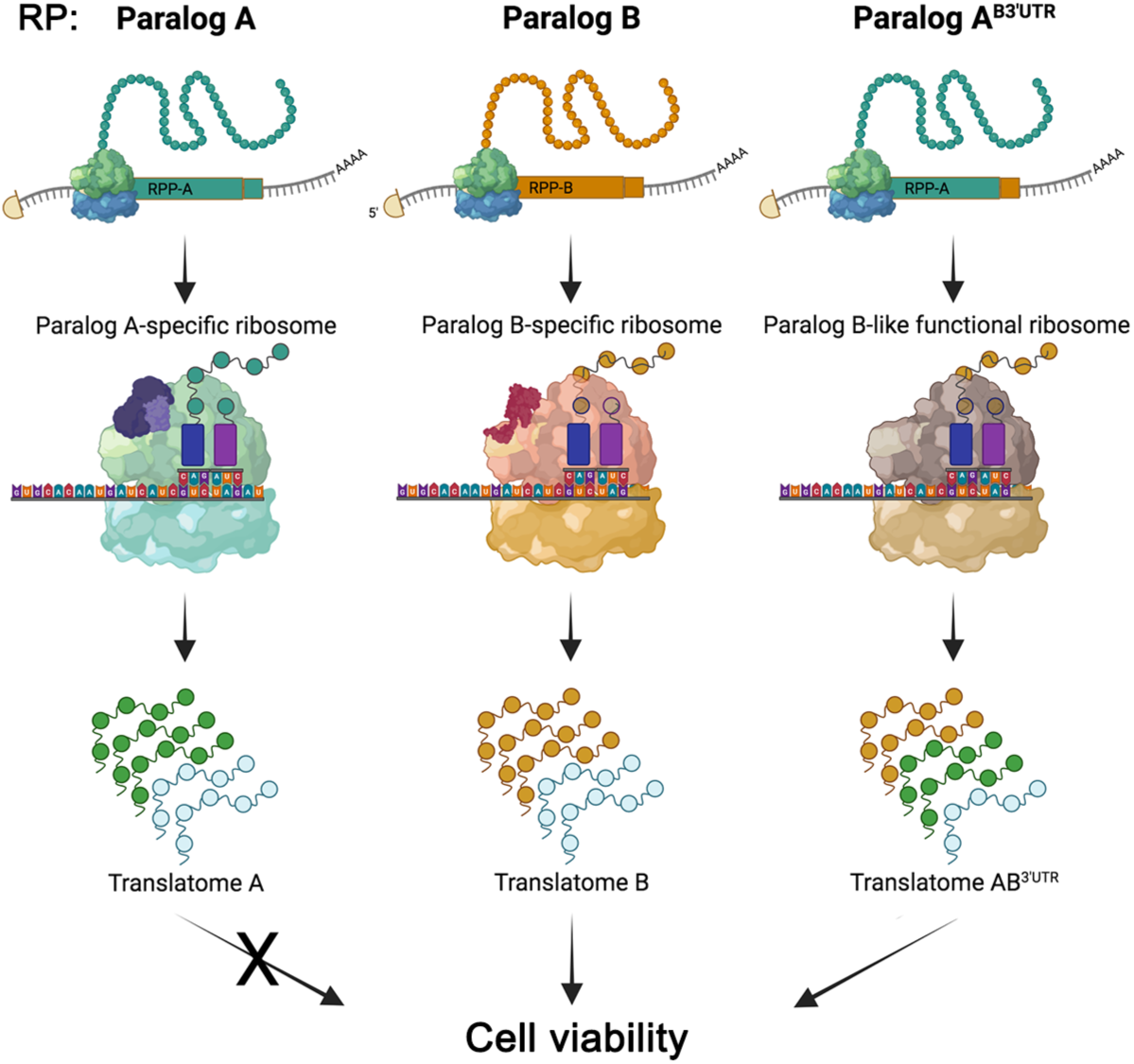

## Introduction

Ribosomes are molecular machines that universally translate proteins and comprise the same basic components: ribosomal RNAs (rRNAs) and ribosomal proteins (RPs). Once considered invariant, recent studies suggest that changes in ribosome composition or modification of any components, such as RPs or specific RP paralogs, rRNA, or ribosome-associated proteins (RAPs), create heterogeneous ribosomes specialized for the preferential translation of specific subsets of mRNAs ^1–6^. Thus, ribosome specialization may play a key role in the regulation of cellular functions, and numerous studies have substantiated its involvement in growth control, stress responses, differentiation, and tissue-specific mRNA translation^2,7–14^. Ribosome modifications and heterogeneity have also been linked to various diseases, including ribosomopathies, neurodegenerative disorders, cancer, and viral infections^15–22^. Moreover, studies made during the COVID-19 pandemic found that the SARS-CoV-2 virus can hijack host ribosomes using its own non-structural protein 1 (NSP1), creating ribosomes tailored to promote viral mRNA translation, while inhibiting host mRNA translation^23–25^. Yet, no comprehensive understanding of ribosome heterogeneity exists for any organism (even under conditions of optimal growth) and, even more importantly, the mechanisms that control ribosome composition leading to specialization remain completely unknown.

To shed light on both the phenomenon and mechanism of ribosome specialization, we performed a systematic analysis of ribosome heterogeneity that arises from differential usage of RP paralogs in the yeast, *S. cerevisiae,* under different growth conditions. In yeast, RP paralogs are gene copies created during gene and chromosome duplication events which lead to 59 RP paralog pairs (*i.e. a* and *b* paralogs) out of a total of 79 RPs present in the ribosome. Given their constitutive expression, paralog gene duplication therefore results in 2^59^ possible combinations of complete ribosomes^26^, although yeast typically contain only around 10^5^ ribosomes^27^. This finding suggests that only a few paralog combinations create the ribosome populations necessary for optimal cellular growth. As ribosome heterogeneity resulting from rRNA or RP modifications can also occur after ribosome biogenesis, heterogeneity that arises from paralog usage alone should be easier to examine. This is because assembled ribosomes are likely to be relatively stable, though ribosome repair pathways might potentially contribute to heterogeneity^28^.

In this study we employed yeast RP paralog deletion libraries and systematic phenotypic screening to identify distinct ribosome populations of specialized function. Another study goal was to elucidate the molecular mechanism that leads to ribosome specialization by examining the translatome and composition of these distinct ribosome populations. As RP paralogs have near identical or identical protein sequences (*i.e.* 95-100% identity), molecular biology tools including immunoprecipitation, immunoblotting, and mass spectrometry of the native paralogs are of limited use. In addition, epitope tags can be used only with a small number of RPs, since in some cases they interfere with integration into the ribosome or ribosome function^29^. However, since paralog gene deletions are not lethal for yeast under optimal growth conditions^30^, we generated single gene deletion libraries for paralogs of both the large and the small ribosomal subunits. We hypothesized that if specific paralogs reside within a single ribosome population, then their individual deletion mutants should exhibit very similar phenotypes. Consequently, phenotypic analysis of the deletion mutants grown under a variety of growth conditions (*e.g.* carbon sources, environmental and proteotoxic stresses) allowed us to cluster ribosome populations and to examine their composition, structure, and translational abilities. We then focused on characterizing ribosomes specialized for growth on oleic acid, which necessitates functional peroxisomes. This approach helped us limit our analysis to a small and well-defined peroxisome-related proteome.

Importantly, we found that such ribosomes confer distinct translatomes and have unique protein compositions that include specific RAPs. In particular, Vps30 (also called Atg6), a component of the yeast phosphoinositide (PI)-3 kinase pathway involved in endosomal protein trafficking and autophagy and an ortholog of mammalian Beclin^31^ is a novel RAP. Vps30 binds to the 40S subunit and is required for the assembly of 80S monosomes when yeast cells are grown on oleate as a carbon source. In addition, we identified the mechanism by which specific paralogs and RAPs associate with RPs to generate specialized ribosomes. We found that switching the 3’-untranslated regions (3’UTRs) between paralog pairs was necessary and sufficient to convert a paralog non-essential for growth under a certain growth condition (*e.g.* growth on oleate or glycerol) to one that is conditionally required. Thus, the 3’UTRs of RP paralog pairs are functional determinants for the association of other cohorts of specific paralogs and RAPs that, together, define a ribosome that confers the efficient translation of select mRNAs. In summary, this comprehensive study aids in defining ribosome heterogeneity in yeast, provides new insights into how specific RP paralogs are assembled into ribosomes, and shows how they recognize and translate different mRNAs to confer optimal cellular growth.

## Results

### RP paralog deletion mutants form phenotypically distinct clusters reflecting paralog-specific ribosome heterogeneity

If specific paralogs are incorporated within a single ribosome population, we hypothesized that their individual deletion mutants should exhibit similar or identical phenotypes that could be clustered according to hierarchy. To test this, we employed near-complete RP paralog gene deletion libraries (*i.e.* comprising 110 out of 135 RPs, including paired and non-paired non-essential RPs) for the individual *a* and *b* paralogs corresponding to the large and small subunits. We next performed a stepwise experimental plan to elucidate the existence and mechanism of action of specialized ribosomal populations (Figure 1). Initially, each of the 110 deletion strains or WT control cells were cultured separately on solid growth media containing different carbon sources (*e.g.* glucose, glycerol, or oleic acid; Figure 2A) at a range of temperatures (*e.g*. 26°C, 30°C and 35°C). In parallel, the individual strains were cultured in liquid growth media under different stress conditions (*e.g*. oxidative, osmotic, or high salt stress; Figure 2B). Solid media was used for measuring growth using the different carbon sources, since growth on oleic acid using liquid media is slow and difficult to assess. Thus, colony size measurement was used as a proxy for growth on the different carbon sources. For assessing cell growth under stress conditions, we compared the growth of deletion strains to wild-type cells on liquid medium, which provides a more direct and precise measurement.

**Figure 1.**
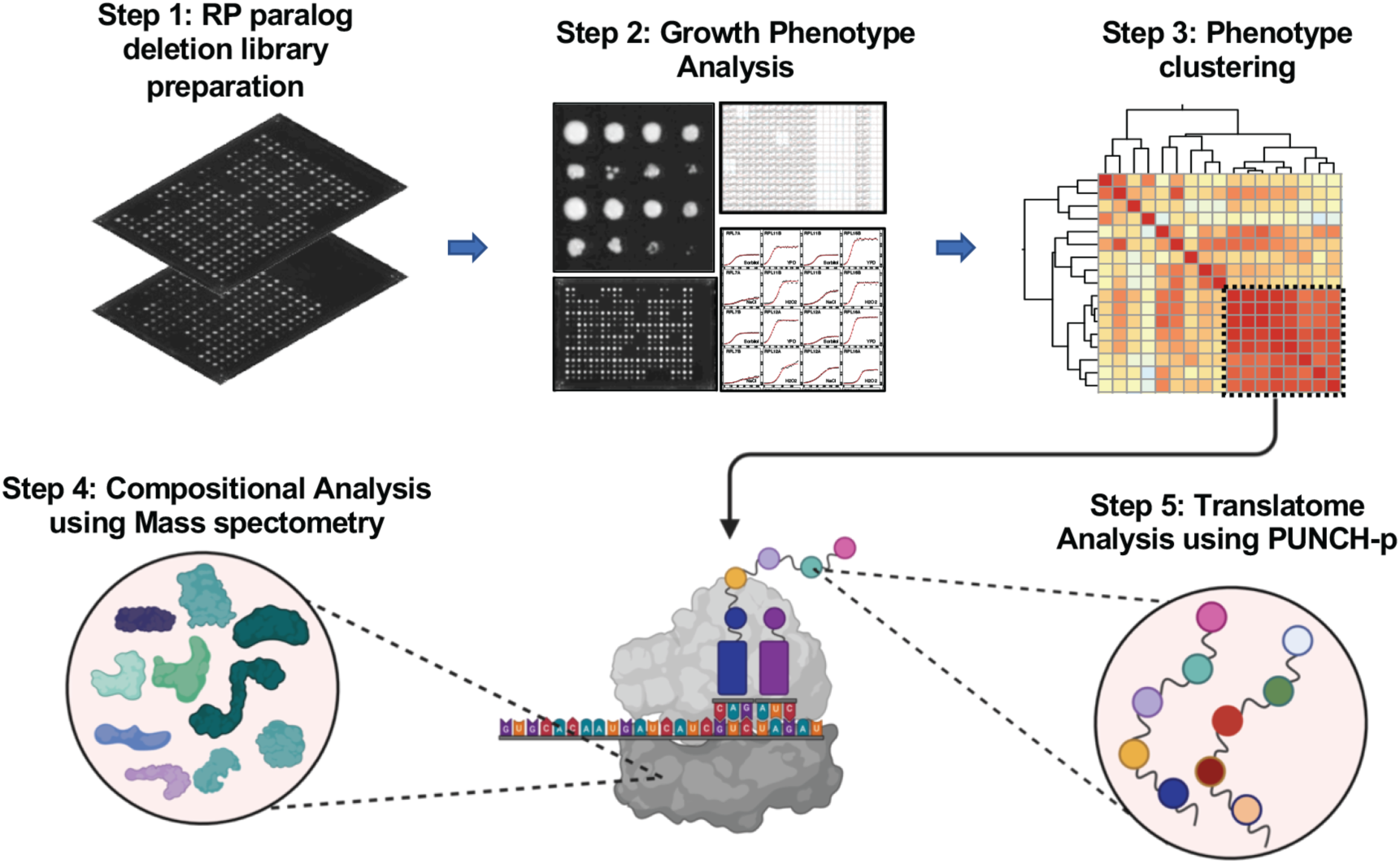
Experimental design to explore specialized ribosomes and their control of yeast cell physiology. Schematic design of the study: Step 1 - creation of deletion libraries of small and large subunit RPs and RP paralogs. Step 2 - growth phenotypes of the deletion strains were measured either using drop tests on solid medium for the different carbon sources (glucose, glycerol, and oleic acid) or in liquid medium for growth under stress conditions (*e.g.* osmotic shock, high salinity, and oxidative stress). Step 3 - cluster analysis of the deletion mutants according to growth phenotype under the different conditions. Step 4-translatome analysis of specific paralog deletion strains using PUNCH-p. Step 5 – determination of the composition of selected paralog-specific ribosomes. Step 6 – resolution of the structure of paralog-specific ribosomes.

**Figure 2.**
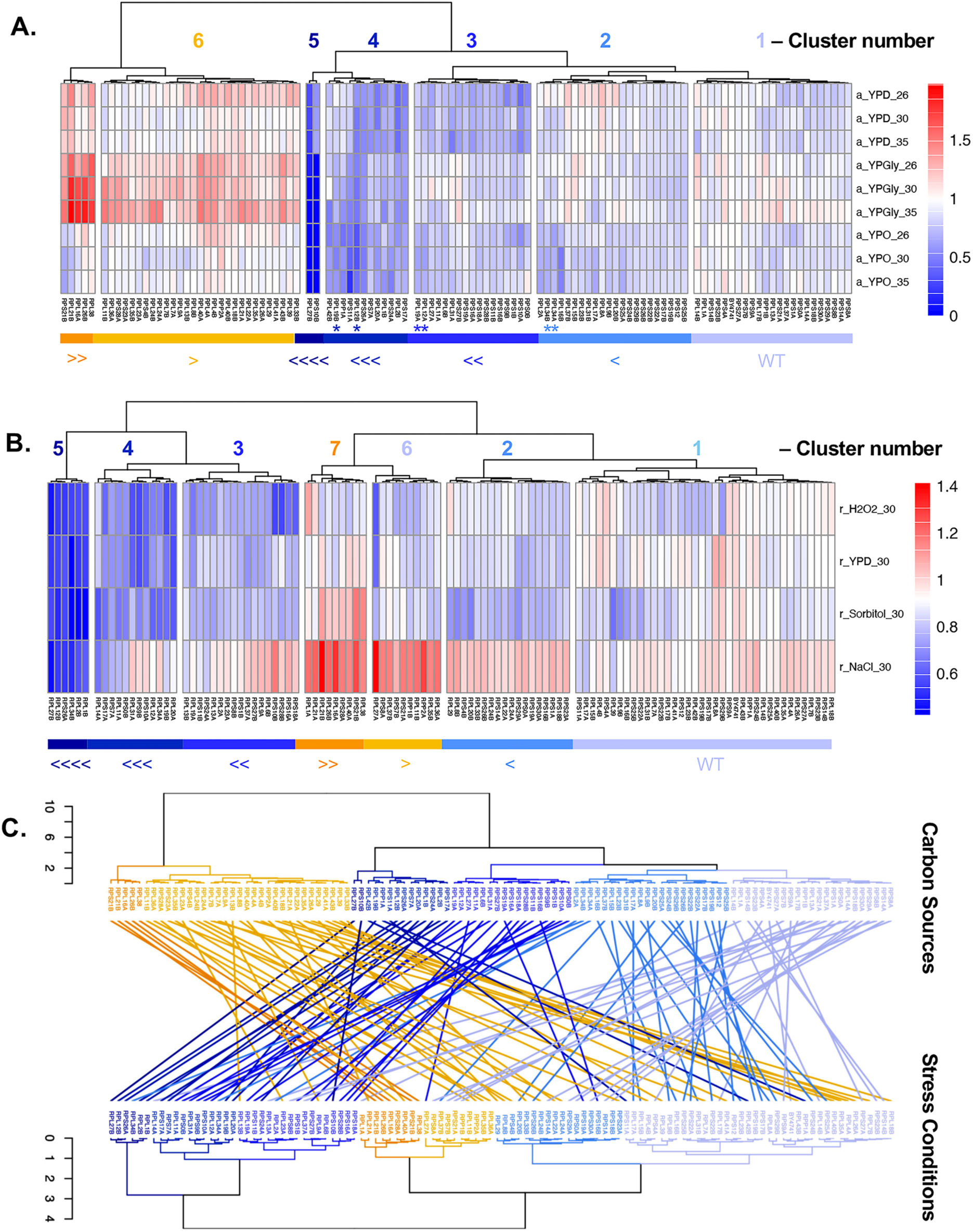
High-throughput screening and cluster analysis of the growth phenotypes of RP deletion mutants (A) A heatmap showing hierarchal clustering of the growth phenotype of RP deletion mutants grown on different carbon sources: Glucose (YPD), glycerol (YPG), and oleic acid (YPO) at 26°C, 30°C and 35°C. Growth was based on colony size measurement. Cluster number is as listed and color-coded as shown in the bar beneath cluster. Strains were classified into seven main clusters relative to the growth of WT cells. Cluster 1-growth equivalent to WT; Cluster 2 – growth similar to WT; Clusters 3-5 – growth much less than WT, in descending order; Cluster 6 – growth better than WT; Cluster 7 – growth much better than WT. Strains marked with stars: *e.g. rpl12bΔ* and *rpl19bΔ* (Cluster 4); *rpl12aΔ* and *rpl19aΔ* (Cluster 3); and *rpl34aΔ* and *rpl34bΔ* (Cluster 2) were selected for further analysis. (B) Hierarchal clustering of the growth phenotype of RP deletion mutants grown under different stress conditions: Control (YPD), Osmotic (1.5M Sorbitol), Oxidative (5mM H_2_O_2_), and High Salt (1.2M NaCl) stress at 30°C. Growth rate was measured and normalized against that of WT cells. As described above, strains were classified into seven main clusters relative to the growth of WT cells. (C) Tanglegram of hierarchal clustering of the growth phenotype in the different carbon sources (top) and stress conditions (bottom). The dendrograms are color-coded based on the cluster type as listed in the heatmaps. Connecting lines are coloured as shown according to the bar in *A* (*i.e.* labeled as in the carbon source map).

To classify paralogs according to phenotypic similarity, we performed Pearson correlation matrix analysis for all growth phenotypes under the different conditions tested. We found that paralog deletion strains form hierarchal phenotypic clusters based on their growth on various carbon sources/temperatures (Figure 2A, Supplementary Figure 1A, and Supplementary Table 1) and under different stress conditions (Figure 2B). Together, 25 out of 49 RP paralog pairs showed anticorrelation or very low correlation (correlation coefficient <0.5) between the pair-wise deletion of either their *a* or *b* partners in conferring growth (Supplementary Figure 1B). Thus, despite the identical or highly similar protein sequences shared among paralog pairs, their individual *a* and *b* deletions tend to show differential growth phenotypes, depending on growth conditions (*e.g. rpl1aΔ* and *rpl1bΔ*, as also shown previously^13^; *rps10aΔ* and *rps10bΔ*, *etc*.) (Supplementary Figure 1B). This indicates that non-pair RP paralogs that phenocopy one another may function together within the same ribosome. Correspondingly, the deletion of paralogs of the same pair tended to yield differential phenotypes, indicating that they belong to different ribosomes having differing capabilities. We also compared the phenotypes of the individual deletion strains between the two sets of conditions to ascertain if the same growth phenotypes observed on the different carbon sources were also observed under stress conditions.

Hierarchal clustering of the growth phenotypes of the paralog deletions grown on different carbon sources (Figure 2A and Supplementary Table 1) defined seven clusters based on growth similarity: (i) little or no growth change compared to WT cells (cluster C1); (ii) slow growth under one condition (cluster C2); (iii) slow growth under two conditions (cluster C3); (iv) slow growth under all conditions (clusters C4 and C5), which we designate as the “core” or functionally dominant cluster given the necessity of these paralogs for growth under all conditions; and (v) faster growth compared to WT cells (clusters C6 and C7). This suggests that paralogs of the core cluster may be ubiquitously present in the general ribosome population or are specific components of ribosomes involved in the translation of mRNAs essential for growth on the different carbon sources tested.

Interestingly, we identified a similar set of phenotypic clusters on stress-conditioned growth (Figure 2B and Supplementary Table 1) and numbered them S1-S7 in the same manner as described for carbon source-based growth. To compare the phenotypic similarity between paralog partners under both sets of growth conditions, we utilized Tanglegram analysis^32^. This showed that functionally dominant clusters, like core clusters C3-C5, were well-preserved between conditions with little rearrangement among the corresponding clusters S3-S5 (Figure 2C). Specifically, 25 out of 31 paralog deletions belonged to these same three clusters and were necessary for robust growth, while only four deletion strains (*e.g. rps11aΔ*, *rpl27aΔ, rpl34bΔ,* and *rpl42bΔ*) grew similar to or better than WT cells under stress conditions, (Figure 2A-C).

We found that 23 out of 34 RP paralogs (68%) belonging to carbon source clusters C6 and C7, which grew better than WT, distributed to stress condition clusters S1, S6, and S7 (*i.e.* 11 strains to cluster S1, 4 strains to cluster S6, and 8 strains to cluster S7), which grow as well as or better than WT cells. This suggests that these paralog deletion strains have a similar level of non-necessity or perhaps some inhibitory function under both sets of growth conditions. In contrast, 24% (*i.e.* 8 out of 34 RP deletion strains) from carbon source clusters C6 and C7 (*e.g. rpl8bΔ, rpl22aΔ, rpl24aΔ, rpl29Δ, rpl33bΔ, rps4bΔ, rps18bΔ,* and *rps23aΔ*) showed impaired growth solely under osmotic stress conditions (cluster S2), implying a stress-specific functional role; while the rest (9%, *i.e.* 3 out of 34 deletion strains; *rpl9aΔ, rpl13bΔ, rpl23aΔ*) grew slower under at least three stress conditions (cluster S3)(Figure 2A-C and Supplementary Table 1). Finally, we found that ∼30% (*i.e.* 13 out of 44; *e.g. rpl2aΔ, rpl13aΔ, rpl14aΔ, rpl34bΔ*, *etc.*) of the RP deletions of clusters C1 and C2 from the different carbon sources redistributed to other clusters under stress conditions (Figure 2C and Supplementary Table 1).

The hierarchical clustering of paralog pairs based on the growth phenotype validated our initial hypothesis and provided valuable insights into phenotypically distinct ribosome populations. It also indicated a high level of functional specificity among paralog pairs, with core paralogs demonstrating functional dominance across all growth conditions, while other paralogs gained or lost functionality depending on the specific growth conditions and cellular requirements.

### Distinct phenotypic clusters appear to delineate functionally distinct ribosome populations

To investigate whether the phenotypically distinct clusters represent different ribosome populations, we examined the genetic interactions between pairs of paralog deletion mutants (Supplementary Figure 2 and Supplementary Table 2). We assessed the growth of RP paralog double deletion strains with the expectation that double deletions within the same cluster (*i.e.* intra-cluster) could be epistatic and yield either no additional defect or perhaps an intermediate effect, as compared to single paralog deletions. This is because we only hinder one ribosome population, whereas deletions between clusters (*i.e.* inter-cluster) might have either a deleterious or rescue effect as we alter two distinct populations of ribosomes. By growing the strains under different growth conditions as in Figure 2B, we found numerous examples whereby inter-cluster double deletions led to growth deficiencies greater than single deletions (*e.g. rpl12aΔ*/*rps26aΔ*, *rps9bΔ/rpl12bΔ,* and *rps24aΔ/rpl12bΔ* strains grew worse, especially on H_2_O_2_-containing medium) (Supplementary Figure 2A). Similar results were seen with cluster S7 (*rpl1aΔ*) where, in combination with clusters S2 (*rpl20bΔ*), S4 (*rps9bΔ*) and S5 (*rps26aΔ*), grew slowly or like the S2, S4, and S5 single deletions alone and lost the >WT growth phenotype typified by cluster S7 strains. In contrast to deleterious growth combinations, we also observed that phenotypically similar clusters S7 and S6 in combination (*rpl1aΔ/ rps9aΔ*) grew better than or equal to their single deletions alone (Supplementary Figure 2A). When we examined several examples of intra-cluster double deletions (Supplementary Figure 2B), we observed that the growth of combinations S5 (*rpl12b/rps26aΔ*) and S3 (*rpl12aΔ*/ *rps31aΔ* and *rpl12a/rps9b* on YPD) were largely unchanged.

Although we note that the growth of S4 combinations *rpl12aΔ* and *rpl31aΔ,* and *rpl12aΔ* and *rps9bΔ*, were less than that of the *rpl12aΔ* strain. Overall, however, these synthetic interactions imply that paralogs belonging to phenotypically distinct clusters likely belong to functionally distinct ribosomes.

### RP paralogs show differential expression during growth

To understand the importance of individual phenotypic clusters and their associated paralogs, we selected growth on oleic acid as a model phenotype. This is because fatty acid metabolism mainly takes place in the peroxisome, which is a small organelle with a limited number of associated proteins induced on oleic acid. Our laboratory previously studied the trafficking of mRNAs encoding peroxisomal proteins in yeast and recent work by others has demonstrated protein translation on yeast peroxisome membranes^33–35^. For further study, we selected the *RPL12B*, *RPL19B*, and *RPL34B* gene paralogs previously identified as being necessary for growth on oleate and for peroxisome biogenesis^36,37^. Along with their cognate partners these paralogs represent three different phenotypic clusters (Figure 2A; clusters C2-4).

As the deletion of *RPL12B* and *RPL19B*, but not that of *RPL12A* or *RPL19A*, led to growth defects on oleate (Figure 2A), we examined by Western blotting the expression levels of the cognate paralogs on both glucose- and oleate-containing growth media at log phase (Supplementary Figure 3A-C). We employed functional genes tagged endogenously with the FLAG epitope upstream of their 3’UTR sequences in the WT background. We first found that Rpl12b is the higher expressed paralog under both conditions, while the Rpl19 and Rpl34 pairs showed similar levels of protein expression (Supplementary Figure 3A-C). Next, we monitored paralog abundance at regular 1.5-hour intervals starting from stationary phase in YPD and observed paralog-specific expression patterns (Figure 3A). We could not perform a similar time-dependent analysis on YPO medium due to the very slow growth rate (*i.e.* avg. doubling time of WT, *rpl12aΔ*, and *rpl1bΔ* cells on oleate was 212, 231, and 264 min, respectively). As described above, Rpl12b-FLAG was expressed at higher levels in the mid-log phase (*e.g.* 1.5-9 hrs) as compared to its cognate paralog, Rpl12a-FLAG, while the levels of both paralogs were similar at either the beginning of the lag (*i.e.* 15 min) or late stationary phases (*i.e.* 9 hrs) (Figure 3A). Since Rpl12b is more abundant at mid-log it may suggest that Rpl12b-containing ribosomes play a central role in that phase, which might explain why the *RPL12B* deletion grows slowly in YPD and YPO (Figures 2A and 3B). In contrast, Rpl19a-FLAG and Rpl19b-FLAG show constant protein expression throughout continuous growth (Figure 3A). Yet, as the *RPL19B* deletion grows slowly (Figure 2A and 3B), this may indicate that its paralog-specific role is independent of protein abundance (Figure 3B). Finally, we observed paralog-specific protein expression for the Rpl34 paralogs, as Rpl34b-FLAG rapidly increases to saturation level in the early log phase (1.5h) (Figure 3A), while Rpl34a-FLAG is lowly expressed at the start and only reaches maximal expression at early stationary phase (4.5h). Thus, there could be paralog-specific roles for Rpl34b-containing ribosomes in the lag phase and perhaps Rpl34a-containing ribosomes in log phase. The paralog-specific expression pattern is consistent with the position in the phenotypic clustering, with paralogs having log-phase functionality showing a stronger phenotype than those with a role in either the lag or stationary phases.

**Figure 3.**
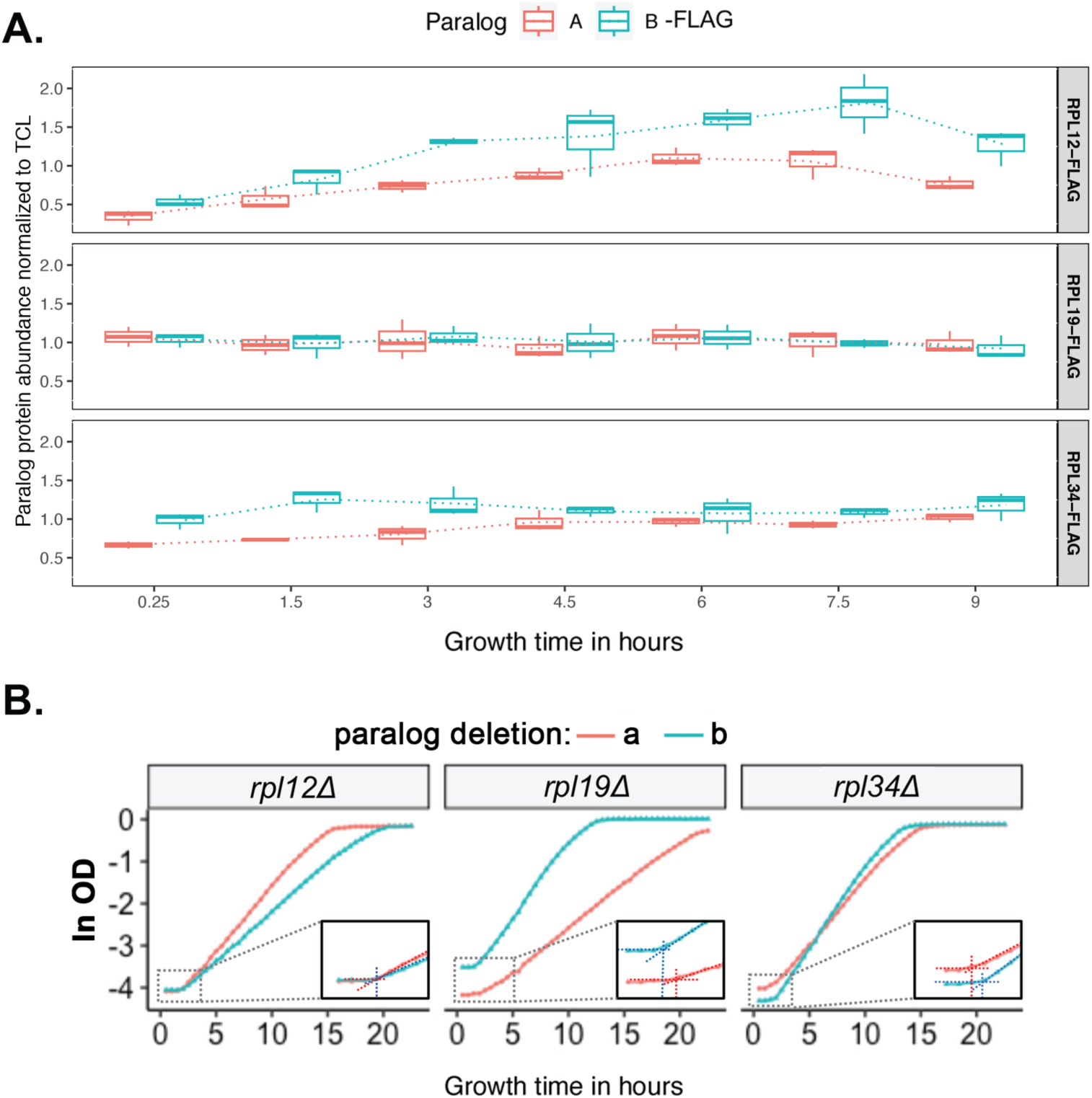
Analysis of RP paralog protein expression and growth on glucose-containing medium (A) Protein abundance of FLAG-tagged RP paralogs as normalized to total cell lysate (N=3 biological repeats). Strains were grown to stationary phase in YPD, then diluted to fresh YPD at O.D._600_ = 0.2 (time 0). Protein extract samples were collected at the indicated time points on the *x* axis. (B) Growth curves of RP paralog deletion pairs in YPD. Deletion strains (*i.e. a* and *b* deletion strains) were grown to stationary phase in YPD, then diluted to fresh YPD at O.D._600_. The O.D._600_ was then measured every 30-40 minutes. *y* axis: ln(O.D._600_). Inset shows the magnified image of the lag phase growth curve with vertical dotted lines to indicate the lag time. Note that mutants *rpl12aΔ* and *rpl19bΔ* grow faster in log-phase as compared to their cognate paralog deletion strains. Cells with *rpl19aΔ* and *rpl34bΔ* have longer lag times than the deletions of their paralog partners.

In parallel, we monitored the growth rate of the individual deletion mutants in glucose-containing YPD medium (Figure 3B). Notably, we observed that the deletion of *RPL12B* resulted in a slower growth rate compared to the *RPL12A* deletion, which correlates with the higher expression of Rpl12b-FLAG protein during mid-log phase (Figure 3A, top panel and Figure 3B, left panel). Moreover, we discovered that the deletion of *RPL19A* led to an extended lag phase and slow growth rate, regardless of the protein expression pattern, which remained consistent in Rpl19a- and b-FLAG cells (Figure 3A and 3B, middle panels). This suggests a potential role for Rpl19a at the start of the growth cycle, although no protein expression differences were observed during non-synchronous cell growth. The deletion of *RPL34B* also caused a delay in the lag phase, which correlates with the increased expression of Rpl34b-FLAG during this phase (Figure 3A, bottom panel; and Figure 3B, right panel).

Since paralog gene deletion could result in changes in the mRNA levels of the remaining paralog and perhaps compensate for the deletion either partially or completely, we examined mRNA abundance of the *a* and *b* paralogs in both wild-type (WT) and deletion backgrounds during growth on YPO and YPD. Our results show reduced expression levels of the *RPL12*, *RPL19*, and *RPL34* mRNAs in the individual paralog deletions, however, more significant reductions were observed in *b* paralog deletion strains on both YPD and YPO. Interestingly, we noted higher expression levels of the remaining paralogs on YPO, perhaps in an attempt to compensate for the deleted paralog. For instance, we observed higher expression of *RPL12* and *RPL19* in *rpl34aΔ* or *rpl34bΔ* deletions (Supplementary Figure 3D and E). Likewise, we observed enhanced *RPL12* and *RPL34* expression in the *rpl19aΔ* or *rpl19bΔ* deletions (Supplementary Figure 3D and E).

### Specialized ribosomes have distinct translatomes

We next investigated whether ribosomes that belong to different clusters have differential translational capabilities, which could explain their different phenotypes. We first attempted Riboseq to determine the translational profile of Rpl12b-versus Rpl12a-containing ribosomes. However, we found that polysomes derived from *rpl12bΔ* cells are RNAase I-sensitive, unlike those derived from *rpl12aΔ* and WT cells, thus, were unable to proceed (data not shown). While the reason behind this selective sensitivity is unclear, it could indicate the existence of structural differences between the two ribosome populations. To overcome this problem, we performed comprehensive translatome profiling using the PUNCH-P technique^38^, which we previously adapted for use with yeast^13^. Isolated nascent polypeptide chains from translating polysome fractions of WT, *rpl12aΔ*, *rpl12bΔ*, *rpl19aΔ*, *rpl19bΔ*, *rpl34aΔ*, and *rpl34bΔ* cells grown on either glucose- or oleate-containing medium were analyzed by mass spectrometry (MS) (Figure 4 and Supplementary Figure 4). We identified nascent peptides corresponding to a total of 755 and 526 proteins, respectively, each detected at least once per biological replica (Supplementary Tables 3-8).

**Figure 4.**
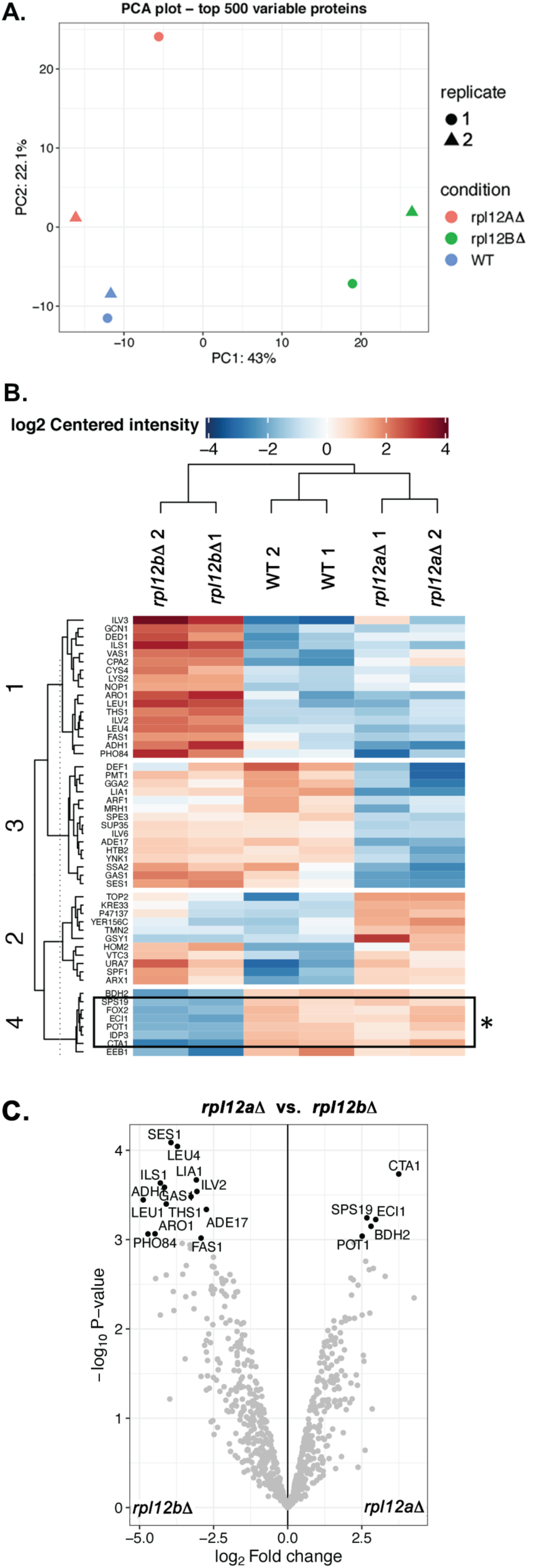
Translatome profiling of the *rpl12△* deletion mutants grown on oleate reveals changes in the translation of peroxisomal proteins (A) Principle component analysis (PCA) of the translatomes of WT, *rpl12a△,* and *rpl12b△* grown on oleate-containing medium (YPO), as analysed using PUNCH-P. (B) Heatmap of hierarchal clustering of nascent polypeptide chains of WT, *rpl12a△,* and *rpl12b△* cells grown in YPO. The *rpl12b△* mutant clusters separately to the *rpl12a△* mutant that co-clusters with WT cells. Translatome cluster numbering is listed as T1-4. Asterisks indicate peroxisomal proteins. (C) Volcano plot of translatomes of *rpl12a△ vs rpl12b△* cells grown on YPO, as measured using PUNCH-P and mass spectrometry.

Principal component analysis and hierarchical clustering revealed that the *rpl12bΔ* translatome in YPO was distinct and clustered separately from that of *rpl12aΔ* and WT cells, which were co-clustered (Figure 4A and B). Importantly, we found that the translation of peroxisome-related proteins (cluster T4) was significantly altered only in *rpl12bΔ* cells on YPO (*i.e.* reduced production of Sps19, Fox2, Eci1, Pot1, Idp3, Cta1, and Eeb1 in oleate-containing medium), while the deletion of *RPL12A* showed almost no effect (Figure 4B and C). In contrast to the effect upon peroxisome-related proteins, a number of proteins show increased translation on oleate upon the deletion of *RPL12B* (cluster T1), including those encoding enzymes involved in branched-chain (Ilv2, Ilv3, Leu1, Leu4) and nucleophilic amino acid synthesis (Lys2, Cys4, Aro1), fatty acid synthesis (Fas1), and proteins that contribute to translation (Gcn1, Cpa2, Ths1, Ils1, Nop2). MEME analysis of these different sets of RNAs whose translatome is altered (*i.e.* downregulated or upregulated) did not reveal the presence of any sequence motifs that might correlate with changes in translation observed on medium containing oleate.

We also found that the translatomes of *rpl19bΔ* and *rpl34bΔ* cells are distinct compared to *rpl19aΔ* and *rpl34aΔ* cells, respectively, in cells grown in YPD and the difference becomes more distinct on YPO medium (see Supplementary Figure 4 and Supplementary Table 5 for *rpl19aΔ* and *rpl19bΔ* cells with WT on YPO, Supplementary Table 6 for *rpl19aΔ* and *rpl19bΔ* cells along with WT on YPD; Supplementary Table 7 for *rpl34aΔ* and *rpl34bΔ* cells including WT on YPD and Supplementary Table 8 for YPO). Like *rpl12bΔ* cells, we observed reduced translation of peroxisome-associated proteins (*e.g.* Pex11, Pcs60, and Vps13) in *rpl19bΔ* cells, as compared to *rpl19aΔ* and WT cells on YPO. Likewise, proteins involved in translation were upregulated under these conditions (*e.g.* Gcn1, Nop4, Efb1).

Since PUNCH-P identifies a fraction of the potential translatome, we determined total protein abundance in the deletion mutants and wild-type cells using label-free MS on whole cell extracts derived from the WT, *rpl12aΔ* and *rpl12bΔ* strains grown either on YPD or YPO (Figure 5, Supplementary Table 11). This analysis revealed the presence of ∼1200-1500 proteins, which was 2-3-fold the level of the PUNCH-P results. Growth on oleate induced substantial changes in protein levels in all three strains, with around two-thirds of proteins differentially expressed compared with YPD cultures (Figure 5A, Supplementary Table 11). As expected, the sets of up- and down-regulated proteins are very similar across all three strains and are enriched for many of the same functional classes (Figure 5D, Supplementary Table 12). For example, GO terms ‘mitochondrion’ and ‘antioxidant’ are enriched among upregulated proteins, while ‘ribosome’ and ‘cytoplasmic translation’ are enriched among downregulated proteins, reflecting the metabolic changes accompanying the needs for growth on oleate. Between strains the scale of changes was smaller for YPD-grown cells (Figure 5B, Supplementary Table 11). Few proteins (*i.e.* <100) were differentially expressed under these conditions between the two *RPL12* deletion mutants. This is reflected in the GO analyses, where most of the enriched terms are shared between the *rpl12aΔ*/WT and *rpl12bΔ*/WT comparisons, and very few enriched functional classes in the *rpl12bΔ*/*rpl12aΔ* comparison (Figure 5E, Supplementary Table 12). The expression profile of *rpl12bΔ* differs substantially from the other strains when grown on oleate (Figure 5C, Supplementary Table 11). Over 250 proteins were differentially expressed between *rpl12bΔ* and *rpl12aΔ*. Most notably, the proteins downregulated in *rpl12bΔ* were enriched for GO terms including ‘oxidoreductase activity’, ‘peroxisomal matrix’ and ‘aerobic respiration’ (Figure 5F, Supplementary Table 12), indicative of a specific defect in the adaptation to oleate and consistent with the strain’s poor growth on YPO. Proteins expressed higher in *rpl12bΔ* included numerous amino acid biosynthesis enzymes, proteasome components, and proteins involved in the response to osmotic stress (Figure 5F, Supplementary Tables 11 and 12). These whole cell proteomics data were highly correlated with the PUNCH-P translatome data (Figure 4). For example, peroxisomal proteins such as Sps19, Fox2, Pot1, and Cta1 were specifically reduced in both the translatome and whole proteome of *rpl12bΔ* cells grown on YPO (Supplementary Tables 4 and 11). Together, these data highlight the impact of the *RPL12B* deletion and suggest that Rpl12b-containing ribosomes are necessary for the translation of specific mRNAs, including those encoding peroxisomal proteins required for growth and survival on oleate.

**Figure 5.**
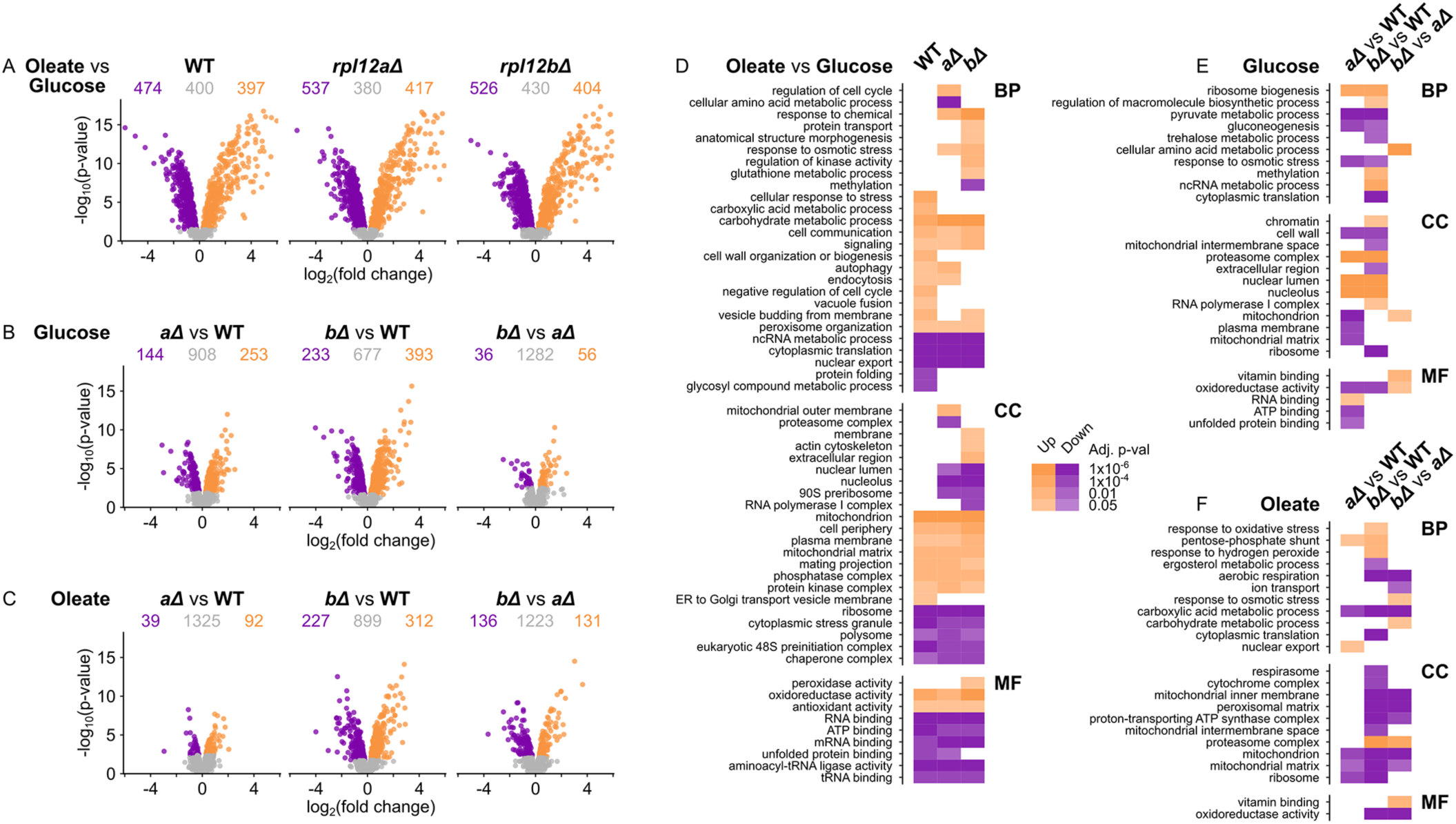
Whole cell proteome profiling of the *rpl12Δ* deletion mutants grown on oleate reveals global changes in protein expression Pairwise comparisons between whole cell proteomes are shown in (A-C). (A) Comparisons between the two growth conditions are shown for each strain. Fold changes are expressed as oleate/glucose. (B & C) Pairwise comparisons between strains for (B) glucose- and (C) oleate-grown cultures. Fold changes are expressed as *rpl12aΔ*/WT (left panel), *rpl12bΔ*/WT (middle) and *rpl12bΔ/rpl12aΔ* (right). For each comparison in (A), (B) and (C), the numbers of differentially expressed (adjusted p-value <0.05) and unchanged proteins (adjusted p-value ≥0.05) are indicated. (D, E & F) GO enrichment analysis of differentially expressed proteins from each comparison. Only enriched terms with adjusted *p*-value <0.05 for at least one comparison are shown. BP – biological process, CC – cellular component, MF – molecular function. Further details are in Supplementary Tables 10 and 11.

### Peroxisome function is altered in RP paralog deletion mutants

Since deletion of *RPL12B* or *RPL19B* inhibits growth on oleate (Figure 2A), as well as the translation of peroxisome-related proteins (Figure 4 and Supplementary Table 11), we examined if the compromised peroxisomal mRNA translation affects peroxisome function. We measured peroxisome induction on oleate, since the peroxisome is the sole site of fatty acid degradation in yeast^39^. To monitor peroxisome induction, we expressed PTS-GFP, a peroxisomal targeting signal fused to GFP, in cells and scored the number of peroxisomes in cells grown on oleate using fluorescence microscopy (Supplementary Figure 4). We found that the number of induced peroxisomes per cell is fewest in *rpl12bΔ* cells, as compared to *rpl12aΔ, rpl19Δ, rpl34Δ* and WT cells (Supplementary Figure 4D). Both *rpl19Δ* (*a* and *b*) mutants were gave similar results, which were less than the other strains, except *rpl12bΔ.* This experiment further supports the idea that peroxisome function is directly affected by the deletion of specific RP paralogs, especially *RPL12B*.

Together, the data suggest that the *rpl12bΔ* and *rpl19bΔ* mutants have an altered translation profile of peroxisome-related proteins that affect their abundance inside the cell and number of peroxisomes, which is a plausible explanation for their slow growth on oleate-containing media (Figures 2A, 4B and C, Supplementary Figure 4B and C). As other findings indicate that mRNAs encoding peroxisomal proteins colocalize with peroxisomes^35^ and that translation can take place on yeast peroxisomes^33^, we investigated the possibility of on-site translation by paralog-specific ribosomes. To accomplish this, we conducted a BioID experiment using strains adapted from Dahan et al. (2022)^33^. Specifically, we deleted individually the *RPL12*A and *RPL12B* paralogs from yeast that express a modified peroxisomal protein (Pex25) fused to BirA and having both Rpl16a and Rpl16b fused to HA and AviTag epitopes, respectively. This technique allowed us to measure the amount of biotinylated AviTag-labeled Rpl16a and b in the *RPL12* paralog-deletion mutants, as compared to cells expressing the paralogs. However, we observed a similar amount of biotinylated AviTag-labeled Rpl16a/b in both the *rpl12aΔ* and *rpl12bΔ* deletion strains, indicating that neither Rpl12 paralog appears to contribute to peroxisome-localized translation in a specific manner (Supplementary Figure 5A and B).

We also examined whether Rpl12 paralogs might show differential association with peroxisomes. To achieve this, we fractionated peroxisomes from WT cells and cells expressing either Rpl12a-FLAG or Rpl12b-FLAG using both sucrose- and subsequent Nycodenz-density gradient centrifugation. These cells also expressed Sec63-RFP to label the endoplasmic reticulum as well as GFP-SKL, which labels peroxisomes. While we successfully isolated a peroxisome-enriched fraction from the cells, as evidenced by both Pex30 and GFP-SKL co-labeling, we were unable to visualize the presence of either paralog (Supplementary Figure 5C). Thus, we were unable to determine whether there is paralog-specific binding, perhaps due to a transient nature of the interaction between ribosomes and peroxisomes.

### Paralogous ribosomes have distinct protein compositions

Next, we directly investigated if the ribosomes formed with each paralog have distinct protein compositions. We isolated 80S monosomes from WT yeast expressing a single C-terminal FLAG-tagged paralog and an untagged paralog and then used anti-FLAG immunoprecipitation to pull down paralog-specific ribosomes from each strain and determined their composition by MS (Figure 6A). Principal component analysis of the MS data revealed that paralog-specific ribosomes which belong to the same phenotypic clusters (*e.g.* Rpl12a and Rpl19a; Rpl12b and Rpl19b; and Rpl34a and Rpl34b; Figure 2A) tend to have similar compositions on both YPD and YPO medium (Figures 6B and C). We plotted the composition of the individual ribosomes on oleate-containing medium (Figure 6D and Supplementary Table 9). The composition of ribosomes in cells grown on glucose-containing medium can be found in Supplementary Table 9. We observed RPs and RAPs that are shared between the paralog pairs (Figure 6D, see dotted line) as well as numerous *a* paralog- and *b* paralog-specific RPs and RAPs (Figure 6D, see distribution along the x and y axes). We detected 371±12 and 222±16 proteins on YPD and YPO, respectively. Specific RAPs like Loc1 and Mrt4 (ribosome biogenesis factors) or Vps30 (a vacuolar PI-3 kinase pathway protein) did not associate with Rpl12a-FLAG-specific ribosomes in YPO. In contrast, Nsa2 (ribosome biogenesis factor), Idp1 (mitochondrial NADP-specific isocitrate dehydrogenase), Fox2 (multifunctional enzyme of the peroxisomal fatty acid beta-oxidation pathway), and others did not associate with Rpl12b-FLAG-specific ribosomes. Some RPs also showed paralog-specific associations, for example despite the similarity to their cognate paralogs both Rpl17a and Rpl22b appear more enriched in Rpl12b-specific ribosomes. Thus, paralog usage in ribosomes help defines their overall proteome.

**Figure 6.**
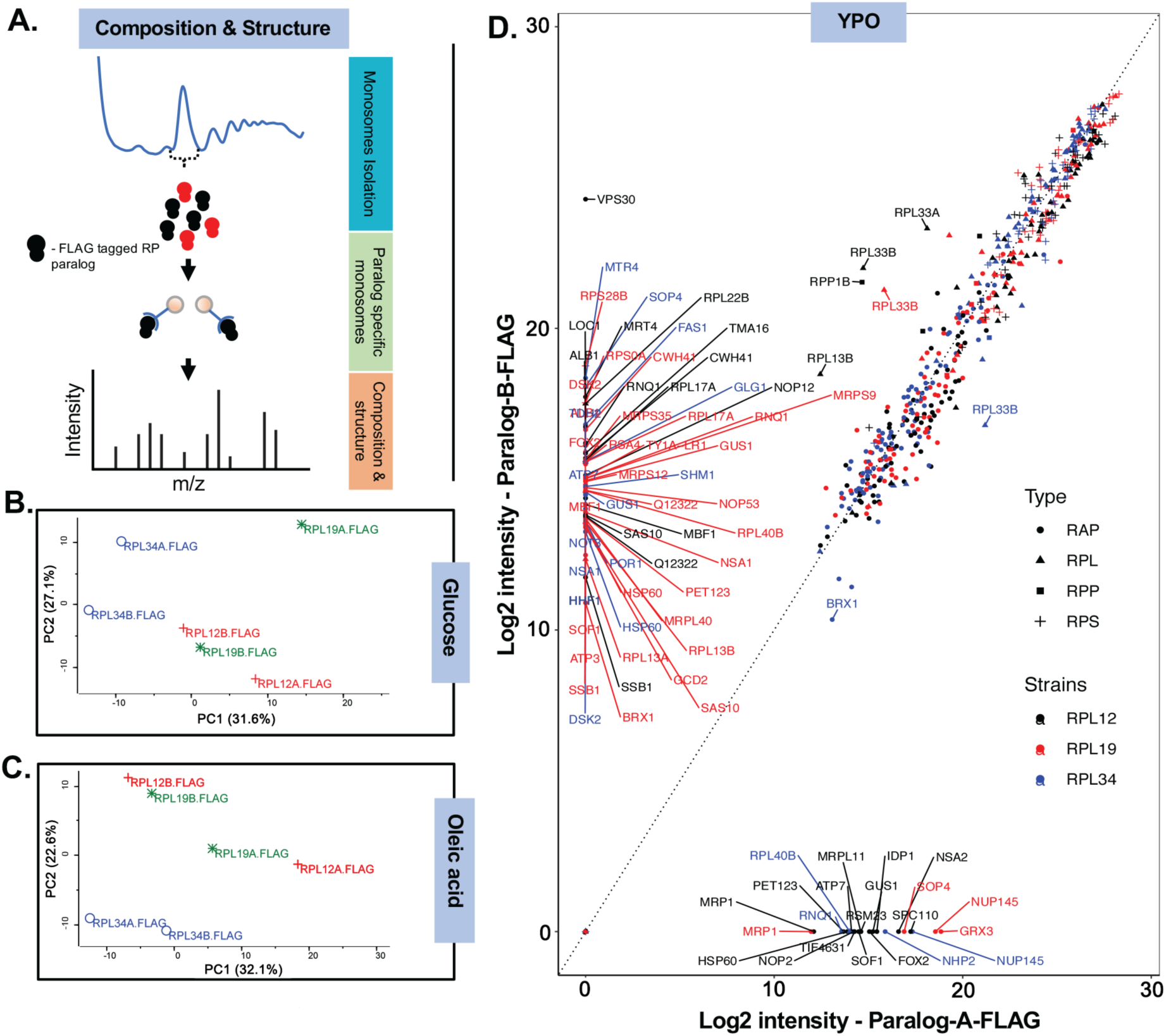
Protein composition analysis of phenotypically distinct ribosomes reveals their heterogeneity (A) Schematic representation of the composition studies. First, total monosomes were obtained from the individual C-terminal FLAG-tagged RP paralog strains by fractionation on sucrose gradients. Next, FLAG-tagged paralog-specific monosomes were affinity-purified using anti-FLAG beads and analyzed using mass spectrometry. (B, C) Principal component analysis (PCA) of the composition of specific FLAG-tagged RP paralog-containing monosomes derived from cells grown on glucose-(YPD) and oleate-containing medium (YPO), respectively. (D) Scatter plots of the quantitative abundance (log2 LFQ) of proteins associated with *a*- and *b*-paralog-specific monosomes. Key: circle - ribosome-associated protein (RAP); triangle - ribosomal large subunit protein (RPL); square - ribosome stalk proteins (RPP); and plus sign - ribosomal small subunit protein (RPS). Color-coded symbols represent the paralog pairs: FLAG-Rpl12a/b (black); FLAG-Rpl19a/b (red); FLAG-Rpl34a/b (blue).

To gain further insight into the ribosome composition, we performed hierarchical clustering of the proteins detected by MS for all paralog-specific ribosomes isolated from both YPD and YPO medium (Supplementary Figure 6A). The YPD and YPO MS data were first analyzed quantitatively and then combined for qualitative clustering to identify similarities and differences in the composition of paralog-specific ribosomes under different growth conditions. Clustering analysis revealed five distinct clusters of detected proteins. Protein composition cluster M1 comprises proteins strongly associated with all ribosomes isolated, which likely indicates their necessity for ribosome function under all growth conditions examined. This cluster comprises 43 RPs, but only one RAP, Stm1, which is involved in translational repression. Composition cluster M2 proteins show a somewhat less unbiased, but likely universal, association of 17 RPs and 35 RAPs with all ribosomes isolated under both growth conditions. In contrast, cluster M3 comprises 7 RPs and 127 RAPs enriched in ribosomes grown in YPD, while cluster M4 comprises 24 RPs and 22 RAPs enriched in ribosomes grown on YPO. Finally, cluster M5 consists of paralog-specific associated proteins, including 8 RPs and 125 RAPs (Supplementary Figure 6A and Supplementary Table 8).

Next, we queried whether the protein composition-specific clusters (Supplementary Figure 7A) form the basis for the phenotypic clusters that result from paralog deletion (Figure 2A). We analyzed similarity and correlation between the phenotypic and compositional clusters using Tanglegram analysis (Supplementary Figure 6B) of the RPs that could be detected by MS. This included 47 RPs, which comprise 43 RPs derived from paralogous pairs and 4 that are unique. We note that the 7 RPs (*e.g.* Rpl6b, Rpl31a, Rps1b, Rps9b, Rps0b, Rps28b, and Rps7a) present in the phenotypic clusters with the most reduced growth (*i.e.* phenotypic clusters C3 and C4, as well as the lesser cluster C3; Figure 2A) largely correlated with composition clusters M1 and M2 under both sets of growth conditions (Supplementary Figure 6B). An exception was Rpp1a, which belonged to composition cluster M4 (Supplementary Figure 6B; deep blue lines). Overall, a complementary relationship was observed between compositional clusters M1 and M2 that corresponded with phenotypic clusters C3 and C4. For example, the paralog-specific composition of cluster M5 correlates with their growth-deficient phenotypes upon deletion, depending on the carbon source. For instance, Rpl17a, which belongs to functionally dominant phenotypic cluster C2 (Figure 2A; poor growth on YPO), has a paralog-specific association with Rpl12b- and Rpl19b-specific ribosomes on oleate. Thus, its deletion phenocopies that of *RPL12B* or *RPL19B* (Supplementary Figure 6B). Likewise, the lower abundance of Rpl29 corresponds to the better growth of yeast under all conditions.

We did observe additional relationships between paralog deletion phenotype and compositional clustering. For example, Rps28b, which belongs to phenotypic cluster C3, did not associate with Rpl19a-FLAG-specific ribosomes under either set of growth conditions (Supplementary Figure 6B). The specific association of Rps28b with Rpl19b-containing ribosomes might suggest a paralog-specific role for Rpl19 during translation, perhaps in anchoring the large and small subunits (Ben-Shem et al., 2011). Unfortunately, some RPs like Rps28a could not be accounted for by compositional analysis due to the lack of unique identifiers, thus, we could not determine whether it fulfills a complementary role under the different conditions. We also noted examples where compositional and phenotypic clustering differed. For example, Rpl36a, like the other 7 RP paralogs present in composition cluster M4, is absent from ribosomes from cells grown on glucose, but present in cells grown on oleate. In contrast, its absence shows either no change or better growth on the different carbon sources. Likewise, while Rps14a is present in compositional cluster M2 (present on both carbon sources), Rps14b is present in compositional cluster M4 (absent in glucose), yet both are present in phenotypic cluster C1 (WT growth). These discrepancies might result from the fact that phenotypic analysis was performed using three growth conditions at different temperatures (26°C, 30°C, and 35°C), while the compositional analysis was conducted under only two growth conditions at 30°C.

Of the 17 paralogs that belong to phenotypic clusters C6 and C7, which show better growth than WT cells, 9 were present in composition clusters M3-4 (Supplementary Figure 6B; yellow lines). Their specific carbon source-dependent association suggests their involvement in growth-dependent cellular functions. In contrast, 7 paralogs from phenotypic clusters C6 and C7 belong to composition clusters M1 and M2, being associated with ribosomes under all conditions (Supplementary Figure 6B). Overall, these results corroborate our idea that the paralogs corresponding to the core phenotypic clusters are either present in most ribosomes or part of functionally dominant ribosomes. Paralogs associated with better growth than WT (*i.e.* cluster C6 and C7) are linked to functionally weaker ribosomes (*i.e.* cluster M2) or exhibit functionality in a growth-dependent manner.

Since the paralog-specific composition of cluster M5 (*e.g.* Rpl17a, Rpl29) effectively explains paralog-specific functionality, we investigated whether paralog-specific RAP interactions contribute to the mechanism. If a RAP contributed to paralog-specific growth, we hypothesized that its deletion would phenocopy that of the deleted paralog. We focused on Rpl12a- and Rpl12b-associated RAPs as the *RPL12b* deletion has strongest effect on growth on oleate and translatome analysis showed reduced translation of peroxisomal proteins. Thus, we individually checked deletion strains encoding the 12 proteins specifically associated with Rpl12b-FLAG monosomes derived from cells grown in YPO (Supplementary Figure 7). We found that the deletion of *NHP6A*, which encodes an HMG1 family member involved in nucleosome remodeling and transcription^40^, grows slowly in both YPD and YPO. In contrast, deletion of its paralog, *NHP6B,* grew like WT cells under both growth conditions (Supplementary Figure 7). Interestingly, we found that Vps30, a component of the yeast PI3-kinase complex involved in protein sorting^41^, is strongly associated with Rpl12b-FLAG ribosomes on YPO. Moreover, the deletion of *VPS30* led to very poor growth on oleate, whereas it grew well on glucose-containing medium (Supplementary Figure 7 and Figure 7A, the latter picture cropped from Supplementary Figure 7).

**Figure 7.**
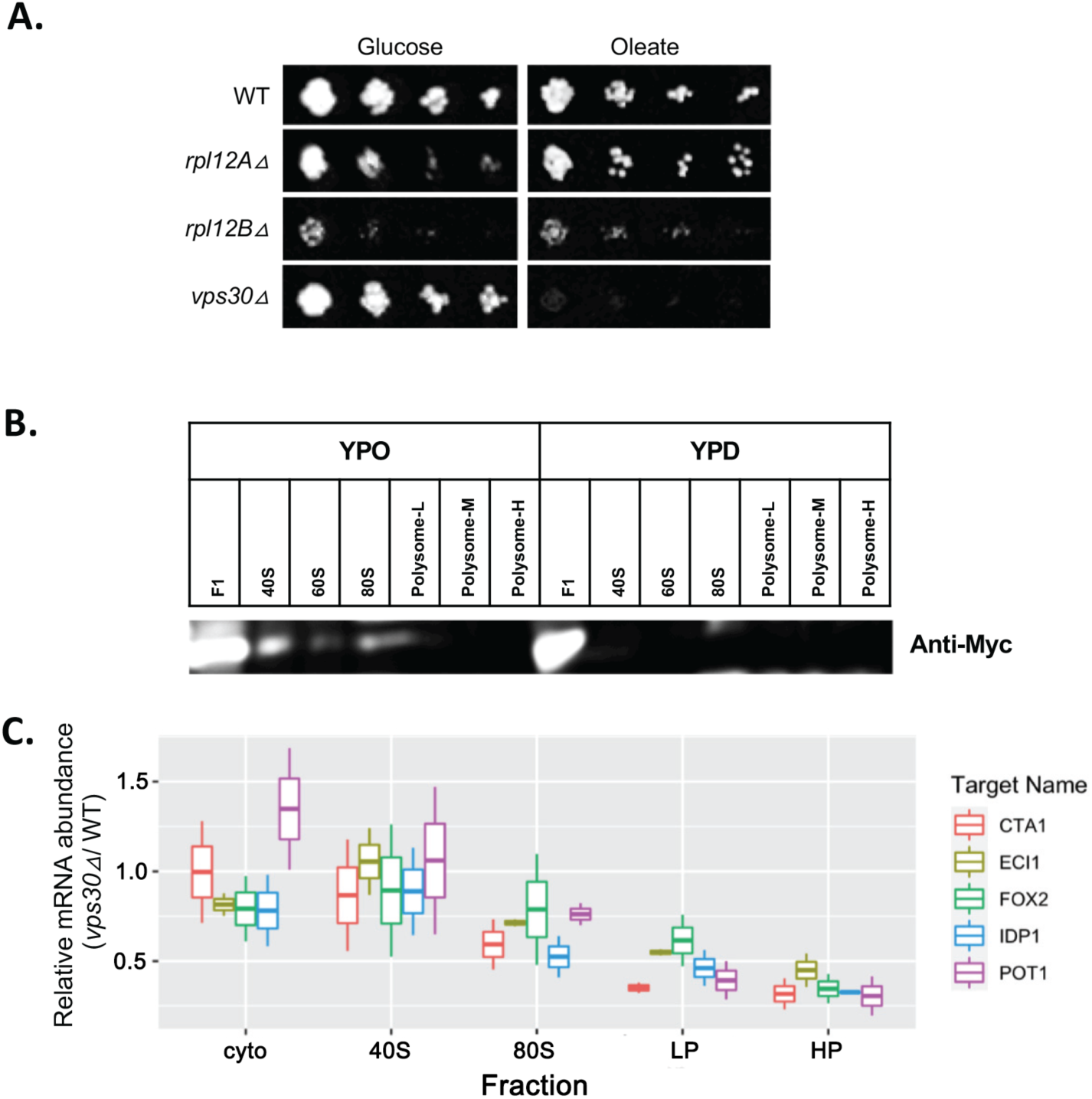
Vps30 binds to the small ribosomal subunit in cells grown on oleate and its depletion causes oleate sensitivity (A) Drop test of WT, *rpl12bΔ, rpl12bΔ*, and *vps30Δ* cells on glucose-(YPD) and oleate-containing medium (YPO). Note that the *vps30Δ* mutants exhibit slower growth only on YPO. (B) Immunoblot of C-terminal Myc-tagged Vps30 cell in different polysomal fractions: cytosol (F1), small subunit (40S), large subunit (80S), light polysomal fraction (L), medium polysomal fraction (M), and heavy polysomal fraction (H) from cells grown on YPO and YPD. Immunoblots probed with anti-Myc antibody. (C) Quantification of the relative abundance of peroxisomal mRNAs in the polysome fractions derived from *vps30Δ* versus WT cells (*e.g.* cytosolic, 40S, 80S, L, M, and H) using sucrose gradient centrifugation and quantification using RT-qPCR analysis. mRNA abundance was normalized to *ACT1* as a housekeeping gene. Cells were grown on YPO medium.

### Vps30 facilitates the formation of paralog-specific 80S monosomes containing peroxisomal protein mRNAs

We investigated how Vps30 interacts with Rpl12b-specific ribosomes by examining its incorporation into the ribosome subunits. We performed polysome fractionation from WT cells expressing MYC-tagged Vps30 and found that it is present in the 40S and 80S fractions, but not the polysome fractions, and only in cells grown on oleate (Figure 7B). Next, we investigated whether Vps30 is involved in the translation of mRNAs encoding peroxisomal proteins. We focused on the mRNAs whose translation was affected by *RPL12B* deletion, as identified by translatome analysis (Figures 4C and D). We measured the abundance of peroxisomal protein mRNAs in the free cytosolic, 40S, 80S, light polysome, medium polysome and heavy polysome fractions by RT-qPCR (as normalized using *ACT1* as a house-keeping control). The relative abundance of these mRNAs (*e.g. CTA1, ECH1, FOX2, IDP1,* and *POT1*) in *vps30Δ* cells was the same as in WT cells in both the cytosolic and 40S (small subunit) fractions, while gradually decreasing from the 80S monosome fraction to the light and heavy polysome fractions (Figure 7C). This suggests Vps30 is required for the efficient formation of the 80S monosomes and polysomes that translate these mRNAs. Since mRNAs are first recognized by the 40S subunit, to which Vps30 binds (Figure 7B), it may act at the rate limiting step of 80S formation.

### Polysome proteomics reveals changes in protein association with translationally active ribosomes in oleate-containing medium and upon Rpl12 paralog loss

We next aimed to identify RAPs more widely associated with polysomes involved in translational regulation specifically during growth on oleate. We analysed the composition of polyribosomes isolated from the WT, *rpl12aΔ*, and *rpl12bΔ* strains grown on glucose- or oleate-containing medium. Samples were crosslinked with formaldehyde, fractions corresponding to monosomes (M), light polysomes (L; 2-4 ribosomes per mRNA) and heavy polysomes (H; five or more ribosomes per mRNA) were isolated, and their compositions analysed using label-free MS. To compare polysome association between conditions or strains, the sucrose gradient fractions were normalized against their respective whole cell extract compositions (Figure 8, Supplementary Figure 8, Supplementary Table 13).

**Figure 8.**
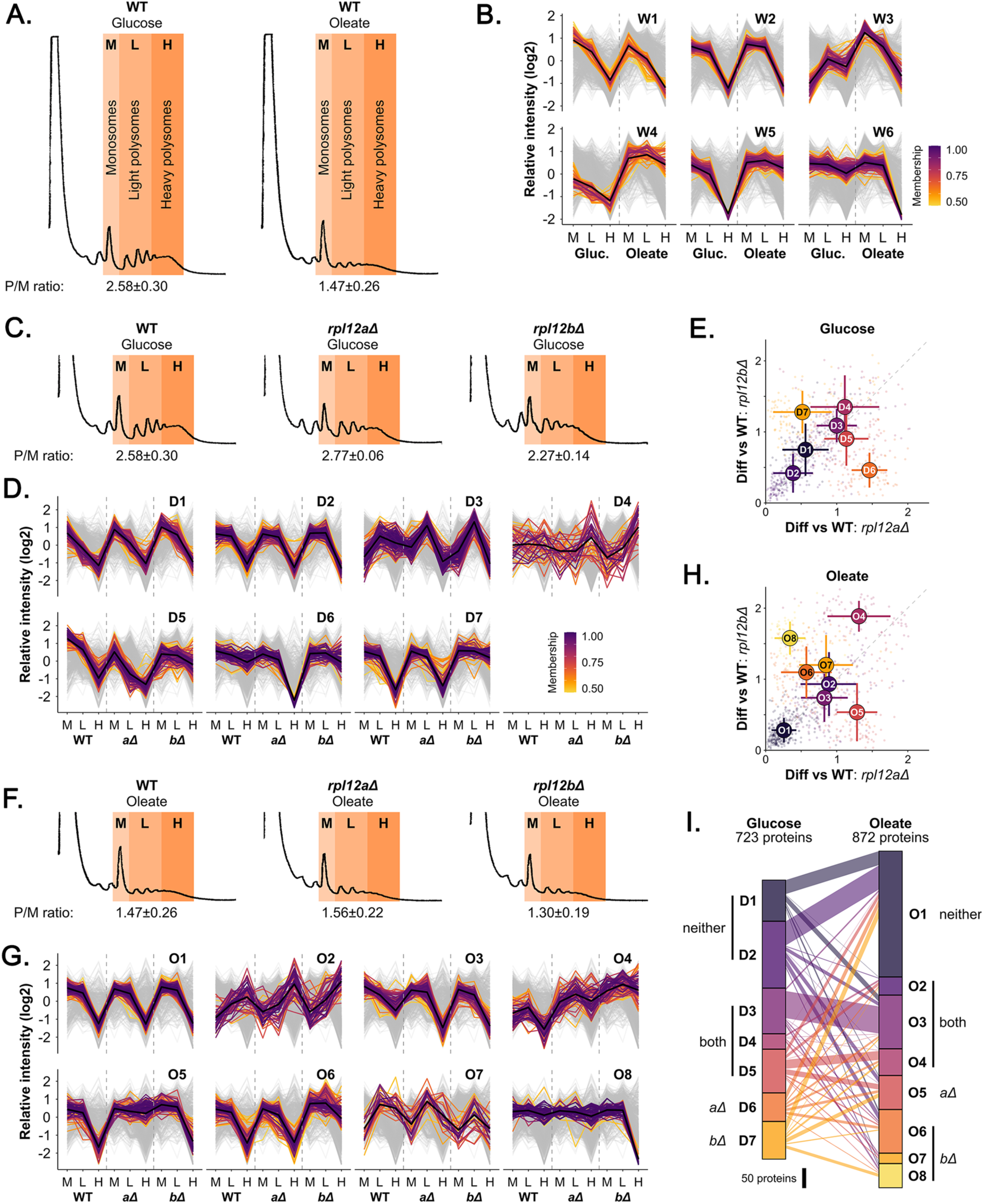
Protein association with polysomes is both media- and Rpl12 isoform-dependent (A) Polysome profiles for the WT strain in glucose- and oleate-grown conditions. The positions of the monosome (M), light polysome (L) and heavy polysome (H) fractions are indicated. The mean polysome-to-monosome (P/M) ratio for each condition is shown underneath (±SD). (B) Fuzzy clustering of normalized protein abundances in WT cells across the sucrose gradient fractions. Proteins with membership values to their respective clusters ≥0.5 are displayed (575 proteins), with all other proteins as the background (grey; 966 proteins). W = WT. (C and E) Polysome profiles for the WT, *rpl12aΔ* and *rpl12bΔ* strains in (C) glucose- and (E) oleate-grown conditions, annotated as in (A). For each strain, the P/M ratio in (E) is significantly lower than its counterpart in (C) (all adjusted p-values <0.001). (D and F) Fuzzy clustering of normalised protein abundances across the sucrose gradient fractions. Proteins with membership values of ≥0.5 to their respective clusters are displayed (glucose: 723 proteins; oleate 872 proteins), with all proteins as the background (grey; glucose: 958 proteins, oleate: 1131 proteins). Polysome profiles and MS data for the WT strain are the same as those presented in (B). (G and H) Comparison of the effects of *RPL12A* and *RPL12B* deletion on polysome association profiles. For each protein, the root mean squared error was calculated between the fractions in the deletion strains and the WT, indicating the difference of the deletion strain profiles from the WT. The mean and SD were calculated for both deletion strains within each cluster and overlaid on the values for individual proteins. (I) Comparison of cluster composition between glucose- and oleate-grown samples. Bars on either side indicate the size of the clusters, links between pairs of clusters represent the number of shared proteins. The general pattern for each cluster is indicated (neither: *rpl12aΔ* and *rpl12bΔ* profiles similar to WT, both: *rpl12aΔ* and *rpl12bΔ* different from WT, *aΔ*: only *rpl12aΔ* different from WT, *bΔ*: only *rpl12aΔ* different from WT).

Proteomic data were reproducible across three replicates (Supplementary Figure 8A and B, Supplementary Table 13). The composition of polysome fractions differed substantially between samples grown on YPD and YPO (Supplementary Figure 8C, Supplementary Table 13). Fuzzy clustering was used to identify sets of proteins with shared polysome association profiles. First, polysomes from the WT strain grown on glucose or oleate were compared to investigate the changes in RAPs under these conditions (Figure 8B). Six major RAP clusters were identified, demonstrating shared patterns of polysome association and how they change between the growth conditions (W1-W6; Figure 8B, Supplementary Table 12). RPs clustered almost exclusively in cluster W3, with high levels observed in both the light and heavy polysome fractions in cells grown on YPD, becoming enriched in monosomes and depleted from polysomes on YPO (Supplementary Table 13, sheets 2 and 5). As expected, the RP profiles match the shapes of the polysome profiles themselves (Figure 8A and B). In contrast, the proteins in cluster W4 show higher levels of relative polysome association in cells grown on oleate (Supplementary Table 13, sheets 2 and 5). Since heavier polysomal complexes are expected to be more translationally active, these proteins might include factors that regulate translation during growth on oleate. Gene ontology (GO) enrichment analysis identified that ‘protein folding’, ‘cellular protein catabolic process’, ‘nucleotide binding’ and ‘small molecule binding’ as descriptive categories enriched among cluster W4 members (Supplementary Table 13). Cluster W5 also shows enrichment in both light and heavy polysomes during growth on oleate-containing medium, as compared with growth on glucose-containing medium. These proteins are a diverse group containing abundant metabolic enzymes many previously shown to bind RNA^42^.

We next characterized the differences in polysome association between the WT and *RPL12* deletion strains. Although there was no significant difference in polysome-to-monosome ratio (P/M ratio) for either deletion strain, as compared to WT (Figure 8C, Supplementary Table 13), our clustering of RAPs reveals strain-specific polysome associations. Seven major clusters of proteins were observed for the polysome association profiles (D1-D7; Figure 8D, Supplementary Table 13). Cluster D1 and D2 profiles are similar for all three strains (Figure 8D and E). Both clusters contain proteins that are enriched in monosomes, variable in light polysomes and depleted from heavy polysomes. They comprise ribosome biogenesis factors, proteasome components, translation initiation and elongation factors, aminoacyl-tRNA synthetases, and chaperones (Supplementary Table 13). Clusters D3 and D4 highlight proteins with shared changes in heavy polysome association in the two *RPL12* deletions (relative to the WT cells), whereby RPs make up nearly 75% of cluster D3, with the remainder comprising RNA-binding proteins (RBPs), chaperones, and translation factors (Figure 8D; Supplementary Table 13). This may indicate a minor reduction in translational activity that was not captured in the global polysome trace (Figure 8C). The proteins in cluster D5 differ from the WT in both deletion mutants, but in different ways (Figure 8D and E). Cluster D5 proteins are retained in heavy polysomes in *rpl12bΔ* cells similar to cluster D7 (Figure 8D). The proteins are mainly involved in the response to oxidative stress and contain glycolytic and other metabolic enzymes. There is also a large overlap in the composition of clusters D5 and D7 (Figure 8D) with cluster W5 (Figure 8B). These RAPs increase their association with polysomes during growth on oleate-containing medium, which suggests a link between the growth defect of *rpl12bΔ* cells and the aberrantly high level of polysome association.

Polysome profiles from oleate-grown cultures showed some inhibition of translation initiation, reflected in significantly lower P/M ratios for all three strains (Figure 8F). However, there were no statistically significant differences, suggesting that bulk translation is not strongly affected by the *RPL12* paralog deletions. Eight clusters were identified in the proteomics data from YPO cultures (Figure 8G, Supplementary Table 13). Clusters O2, O3 and O4 represent proteins with shared changes in polysome association in the *rpl12aΔ* and *rpl12bΔ* deletion strains compared with WT (Figure 8G and H), with clusters O2 and O4 showing enrichment in heavy polysomes and cluster O3 instead showing depletion (Figure 8G). The proteins in clusters O6, O7 and O8 show *rpl12bΔ*-specific polysome association profiles (Figure 8G and H). Clusters O7 and O8 are depleted from heavy polysomes in *rpl12bΔ* cells, suggesting they might contain proteins that positively regulate translation during growth on oleate, which is defective in this strain (Figure 2A). Few GO terms are enriched in these clusters, but their members include well-characterized RBPs, like Puf3, Scp160, Scd6 and Sbp1 (Supplementary Table 13)^43–45^. Proteins in cluster O6 might also contribute to translational regulation in the opposite manner. These are predominantly enriched for mitochondrial enzymes involved in aerobic respiration.

The two sets of clusters were then compared to investigate whether the patterns of polysome association are similar independent of growth conditions (Figure 8I). We found a substantial overlap between clusters which contain proteins that do not differ from WT in either *RPL12* paralog deletion strain (*e.g.* clusters D1 and D2 overlaps with cluster O1). Additionally, clusters D3 and O3 overlap very strongly (Figure 8I). These clusters contain proteins for which polysome association in both deletion mutants differs from WT and includes most RPs. However, there were no strong overlaps for the paralog-specific clusters (*e.g.* D6-D7 and O5-O8) with clusters from the opposite growth condition (Figure 8I). Paralog-specific changes in polysome association profiles are therefore highly context-dependent, with different sets of proteins changing between the strains during growth in glucose and oleate. Taken together these analyses show that growth on oleate changes the ribosome-interaction patterns of a wide range of proteins. It also shows that loss of either *RPL12* paralog results in multiple paralog-specific differences with WT cells grown on either glucose- or oleate-containing medium. Ultimately, *rpl12bΔ*-specific changes in polysome composition during oleate growth are likely to contribute to the differences in protein synthesis and consequently total protein levels that we observed (Figures 4 and 5), which themselves result in condition-specific growth defects.

### Paralog-specific 3’UTR sequences control the formation of specialized ribosomes

Our experiments regarding paralog-specific ribosomes raised a key question, *i.e.* how do cognate paralogs having the same coding sequence become incorporated into composition-specific ribosomes? As the 3’UTRs of the mRNAs encoding paralog pairs are unique, we hypothesized that switching the UTRs between cognate paralogs might switch their roles in cellular function. This could be the case if the 3’UTRs help recruit proteins (*e.g.* RAPs or specific RPs) during paralog translation and then incorporate them into a complex with the growing paralog polypeptide that undergoes assembly into a functional ribosome.

To answer this question, we replaced the unique 3’UTR of *RPL12A* with the 3’UTR of *RPL12B* (Figure 9A) in the genome and then deleted *RPL12B* to test whether the chimeric *RPL12A-B^3’UTR^*could rescue the *rpl12bΔ* deletion phenotype as the sole paralog expressed. Importantly, this genetic modification successfully complemented the absence of *RPL12B* when cells were grown on oleate-containing medium at 37°C (Figure 9B). Moreover, we found that the rescue mediated by endogenous *RPL12A-B^3’UTR^* was fully identical to that observed upon episomal overexpression of either *RPL12A* or *RPL12B* in *rpl12bΔ* cells (Figure 9B). Thus, while *RPL12A* paralog can rescue *rpl12bΔ* cells, it is only upon non-native overexpression. In contrast the *RPL12A-B^3’UTR^* allele rescues at the endogenous level of *RPL12A* expression in the absence of *RPL12B*, whereas native *RPL12A* expression alone cannot. Thus, gene dosage cannot account for the rescue by the *RPL12A-B^3’UTR^*allele.

**Figure 9.**
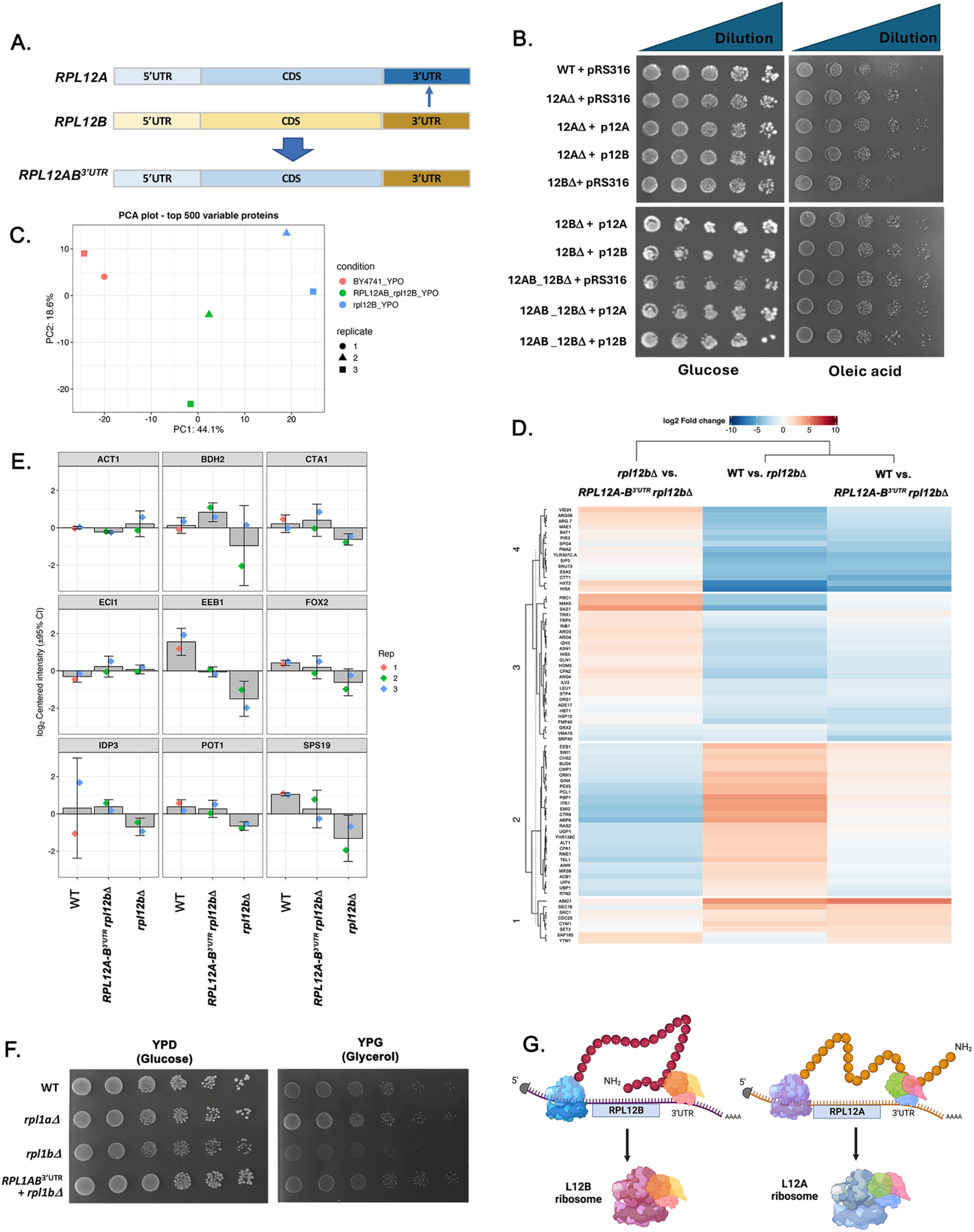
3’UTRs define paralog identity: Substitution of the 3’UTR of *RPL12A* with the *RPL12B* 3’UTR rescues the growth phenotype of *rpl12bΔ* cells on oleate (A) An illustration of the 3’UTR switch made between paralogs. The 3’UTR of *RPL12A* is replaced with that the 3’UTR of *RPL12B*. (B) Drop test of WT, *rpl12a*Δ (12AΔ), *rpl12b*Δ (12BΔ) and *RPL12AB^3’UTR^ rpl12bΔ* (12AB_12BΔ) cells transformed either with a control plasmid, pRS316 or with centrosomal plasmids expressing *RPL12A* (p12A) or *RPL12B* (p12B) under the control of their native promoters. Cells were grown either on YPD (left) or YPO (right) plates at 37°C. Images were taken after 2 days for cells grown on YPD and 4 days for cells grown on YPO. (C) PCA analysis of the top 500 variable proteins identified in the PUNCH-P test of WT, *rpl12b*Δ and *RPL12AB^3’UTR^ rpl12bΔ* cells grown on oleate-containing medium (YPO). (D) Heatmap of hierarchal clustering of nascent polypeptide chains of WT, *rpl12b△*, and *RPL12AB^3’UTR^ rpl12bΔ* cells cells grown in YPO. The *RPL12AB^3’UTR^ rpl12bΔ* chimera clusters separately to the *rpl12b*Δ mutant. (E) PUNCH-P measurements of peroxisomal and peroxisome function-associated proteins derived from PUNCH-P analysis in (D). The histograms represent the relative levels of the proteins with the bars representing 95% confidence interval (CI). Act1 serves as a representative non-peroxisomal protein. (F) Substitution of the 3’UTR of *RPL1A* with the 3’UTR of *RPL1B* confers the growth of *rpl1bΔ* cells on glycerol. Drop test of WT, *rpl1a*Δ, *rpl1b*Δ and *RPL1AB^3’UTR^ rpl1bΔ* cells grown on YPD (left) and YPO (right) plates at 30°C. Images were taken after 3 days for cells grown on YPD and 4 days for cells grown on YPO. (G) A model for the translational association of different RAPs with specific RP paralogs. During the translation of *RPL12B* (left) or *RPL12A* (right), RAPs that associate with the paralog-specific 3’UTRs interact with the nascent polypeptide. This association allows both the paralog and associated RAPs to then assemble into composition-specific ribosomes that preferentially translate mRNA subsets.

Since increased levels in gene expression of the chimeric allele might facilitate rescue, we examined the levels of *RPL12* RNA in WT, *rpl12bΔ*, and *RPL12A-B^3’UTR^ rpl12bΔ* cells by RT-PCR. However, we found no significant change in *RPL12* mRNA levels between *rpl12bΔ* and *RPL12A-B^3’UTR^ rpl12bΔ* cells, which were essentially half that of WT levels. Overall, these results suggest that *a/b* paralog 3’UTR swapping creates functionally distinct ribosomes, *i.e. RPL12A-B^3’UTR^*behaves more like *RPL12B* than *RPL12A*.

To verify this idea, we performed PUNCH-P on WT, *rpl12bΔ,* and *RPL12A-B^3’UTR^* cells grown on oleate-containing medium. PCA analysis (Figure 9C) revealed that all three strains gave non-identical results, with the MS data (Figure 9D) reflecting the intermediate nature of the *RPL12A-B^3’UTR^* cells, *i.e.* being neither identical to either WT or *rpl12bΔ* cells. We specifically examined the levels of peroxisome-associated proteins shown previously to be downregulated in *rpl12bΔ* cells (Figure 4B-C) and observed an intermediate level of these proteins in *RPL12A-B^3’UTR^* cells, relative to WT or *rpl12bΔ* cells (Figure 9E). Notably, these same proteins (*e.g.* Bdh2, Cta1, Eeb1, Fox2, Idp3, Pot1, and Sps19) were largely reduced in *rpl12bΔ* cells, as seen previously (Figure 4B-C), but were expressed higher in *RPL12A-B^3’UTR^* cells. Thus, 3’UTR switching appears to confer partial Rpl12b functions to Rpl12a in *rpl12bΔ* cells.

To further verify that 3’UTR switching creates functionally distinct ribosomes we examined whether swapping the 3’UTR of *RPL1A* with that of *RPL1B* could allow cells to grow well on glycerol-containing medium (Figure 9F), which we previously demonstrated to result from specialized ribosomes containing Rpl1b^13^. We found that *rpl1bΔ* cells expressing the *RPL1A-B^3’UTR^* chimera grew much better on glycerol-containing medium than *rpl1bΔ* cells and were more like WT cells. Thus, the distinct 3’UTRs of RP paralogs appear to confer the formation of specialized ribosomes in yeast.

## Conclusions

Recent works have illustrated the behavior of cognate RP paralogs to generate ribosomes specialized for function^1–6^. However, these findings were derived from specific paralog pairs or isoforms in different organisms, while the complete orchestration of RP paralogs that create heterogeneous ribosomal population has not been elucidated for any organism. Given the plethora of conditions under which yeast can live, we have employed them as a model to study ribosome heterogeneity. We took advantage of the fact that single RP gene deletions are viable and hypothesized that the deletions corresponding to a given population of ribosomes would yield the same phenotype. We clustered the paralogs according to phenotype similarity by deletion library screening using a variety of different growth conditions (*e.g.* carbon source, environmental and proteotoxic stresses) (Figures 2A-C). By examining synthetic interactions between paralog deletion strains we verified that the phenotypic clusters likely represent distinct ribosome populations and found that inter-cluster double deletions tend to have synthetic deleterious or suppressor effects, whereas deletions within a cluster have lesser effects (Supplementary Figure 2).

We selected three pairs of paralogs (*e.g. RPL12, RPL19*, and *RPL34*) representing different phenotypic clusters to study their different responses on oleic acid, which requires peroxisome function, as well as their composition and translatomics. We first examined protein levels of the paralogs and found that their expression is specific to some degree during the different phases of growth (*e.g.* lag, log, and diauxic shift). For example, Rpl12b is highly expressed during logarithmic phase, while Rpl34b is expressed faster during lag phase and reaches a saturation in early log, as compared to their cognate partners (Figure 3). This may indicate the existence of temporal requirements for the different paralogs during growth, which could help form the basis for phenotypic clustering (*i.e.* phase-specific paralog expression could define specific ribosomes). We previously demonstrated that episomal expression of one paralog can complement the loss of the other in the case of *RPL1* and *RPL2*^13^, and here for *RPL12* (Figure 9B). However, genomic levels of paralog expression do not complement the loss of *RPL12B* and *RPL19B*, and the other pairs grouped separately by hierarchal clustering (Figure 2A and B). Moreover, changes in *RPL1A* and *RPL2A* mRNA and protein levels observed upon the deletion of their *b* paralogs could not complement deletion phenotype^13^. To rule out this possibility for the *b* paralog deletions of *RPL12, RPL19*, and *RPL34*, we examined the deletion strains for changes in gene/protein expression that might disguise/complement the deletion. However, we did not observe compensatory changes in RNA and protein levels that might compromise the observed phenotypes (Supplementary Figure 3).

Given their differential phenotypic clustering and sensitivity to growth on different carbon sources, we studied the translatomes of the *RPL12, RPL19*, and *RPL34* paralog deletion using PUNCH-P (Figure 4, Supplementary Tables 2-7). Interestingly, we found that paralog usage produces ribosomes specialized for growth on oleate, which necessitates peroxisome function. We found that the translation of a number of peroxisomal and peroxisome-related proteins is greatly reduced in *rpl12bΔ* and *rpl19bΔ* mutants, which likely explains their slow growth phenotype in oleic acid-containing media (Figure 4) and reduced number of peroxisomes (Supplementary Figure 4D). Although we sampled a relatively small fraction of the entire potential proteome using PUNCH-p (∼10-15%), it was notable that proteins connected to peroxisome-function were most strongly affected especially in the *rpl12bΔ* cells, in which we also found that the level of peroxisome induction to be low (Supplementary Figure 4D). Whole cell proteome analysis of WT, *rpl12aΔ* and *rpl12bΔ* cells grown on glucose- and oleate-containing medium (Figure 5, Supplementary Table 11), which measured the steady state levels of 2-3-fold more proteins than PUNCH-P, revealed that >250 and >500 proteins are differentially expressed between *rpl12bΔ* and *rpl12aΔ* cells, and *rpl12bΔ* and WT cells, respectively, on YPO (Figure 5C). This also included proteins directly involved in peroxisome function, which correlates with the PUNCH-P data and illustrates the necessity of Rpl12b-containing ribosomes for growth on oleate (Figure 5F, Supplementary Table 12).

Although we investigated the possibility of localized translation by paralog-specific ribosomes on peroxisomes, we did not find conclusive evidence for it (Supplementary Figure 5). Thus, to better understand how *a* and *b* paralog pair-containing ribosomes differ in function, we investigated their composition by pulldowns of ribosomes containing the different FLAG-tagged forms of Rpl12, Rpl19, and Rpl34 grown on either glucose- or oleic acid-containing medium and MS (Figure 6). Importantly, we found that monosome composition is highly distinctive, depending upon the paralog incorporated, and that the differences derive primarily due from the association of specific RAPs and in some cases other RP paralogs (Figure 6D and 7A). As with the phenotypic screening and translatome/proteome analyses, the results from the compositional study strongly support the idea that specialized ribosomes are distinct from those less specialized for growth under certain conditions (*e.g.* Rpl12b-containing monosomes are distinct from Rpl12a-containing ribosomes and recruit RAPs necessary for the efficient translation of peroxisomal proteins). Thus, the basis for ribosome specialization is primarily connected to paralog selection.

Since RAPs appear responsible for compositional differences, we investigated whether the deletion of specifically associated RAPs has the same phenotype as paralog deletion. Importantly, we found that deletion of *VPS30*, which is specifically bound to Rpl12b-FLAG-specific ribosomes in oleate-containing growth conditions, resulted in the same slow growth as *RPL12B* paralog deletion (Figure 7A). As we later found that Vps30 first binds to the 40S small subunit (Figure 7B), it could be that it helps Rpl12b-specific large subunits to recognize peroxisomal mRNAs bound to the 40S and thus assemble into monosomes under these conditions. Correspondingly, the deletion of *VPS30*, like that of *RPL12B*, decreases the translation of peroxisome-related mRNAs (Figure 7C) and lowers their levels in the polysome fraction, in which the rate limiting step is 80S monosome formation. Overall, ribosome composition analysis suggests that the interaction of RAPs may be responsible for ribosome heterogeneity, perhaps by guiding the small subunit of the ribosome to select specific mRNAs and interact with paralog-specific large subunits for specialized translation. In the case of Rpl12b-containing ribosomes, we suspect that Vps30 works as a gatekeeper for the efficient recruitment and translation of select mRNAs, *i.e.* those encoding peroxisomal proteins. Thus, it could function like the SARS-CoV-2 coronavirus protein, NSP1, which binds to host ribosomes near the mRNA entry tunnel, to inhibit host mRNA translation, and promote efficient viral mRNA translation^23,25^. More work is required to determine how Vps30 specifically fulfills its role as a RAP in yeast cells grown on oleate, but not on glucose. Likewise, it is unknown whether mammalian Vps30 (Beclin), which is associated with autophagy-related brain disorders and neurodegenerative disease, has a similar function.

To better understand the level of changes associated with translationally active ribosomes, we measured polysome proteomics associated with the deletions of the *RPL12* paralogs on both growth media, as compared to WT cells (Figure 8, Supplementary Figure 7, and Supplementary Table 13). We observed substantial changes in the composition of polysome fractions between samples grown on YPD and YPO and clustering revealed sets of proteins with shared polysome association profiles and how they changed between the growth conditions. We found that paralog-specific changes in protein association with the monosome and polysome fractions were context-dependent, with different sets of proteins changing between the strains during growth in glucose and oleate. The deletion of either *RPL12* paralog resulted in multiple paralog-specific differences with WT cells. That said, we observed *rpl12bΔ*-specific changes in polysome composition during oleate growth (*e.g.* clusters O7 and O8, that contain proteins involved in translational control; Figure 8G) that likely contribute to the observed differences in protein synthesis and total protein levels (Figures 4 and 5), resulting in condition-specific growth defects.

Finally, a central unanswered question is how specialized ribosomes are created using paralog pairs that have near, if not full, identity at the coding level. Since the 3’UTRs are distinct, we reasoned that the differences in ribosome composition might arise during translation, in which the paralog pair-specific 3’UTRs recruit different RAPs or other RP paralogs to interact with the translating paralog to create a paralog-specific protein complex. Work by Mayr and colleagues has already demonstrated that during translation the 3’UTR can recruit select proteins to the nascent polypeptide chain and, thus, allow for the formation of specific protein complexes^46^. We hypothesized that this would be a convenient mechanism to control protein-protein interactions between RAPs and RPs that could lead to the formation of specialized ribosomes upon final assembly. Importantly, we demonstrated 3’UTR switching between *RPL12B* and *RPL12A* (Figure 9A) successfully led to role switching between the paralogs on oleate-containing medium, rendering *RPL12B* amenable to deletion, but without leading to the same growth defect as observed with *rpl12bΔ* cells on oleate (Figure 9B). PUNCH-P translatome data for the *RPL12AB^3’UTR^ rpl12bΔ* cells revealed an intermediate effect upon translation, which likely explains their enhanced growth on oleate relative to *rpl12bΔ* cells (Figure 9D and E). This included proteins necessary for peroxisome function (Figure 9E), hence it appears that the *RPL12B* 3’UTR is largely sufficient to confer specialized ribosome functions under these conditions. Importantly, we could show a similar 3’UTR requirement for *RPL1A* and *RPL1B* role switching in order to confer growth on glycerol (Figure 9F), which necessitates the paralog-specific translation of RNAs encoding nuclear-encoded mitochondrial proteins^13^. Together, our results suggests that RP paralog pairs employ similar protein sequences, likely to maintain the core composition and structure of the ribosome, while their unique UTRs allows them to interact specifically with other proteins during translation to lead to the creation of specialized ribosomes (see model, Figure 9G). We recognize then that this mechanism for the creation of specialized ribosomes could be common to all eukaryotes. Future studies will be necessary to characterize and determine the extent of other specialized ribosomes in yeast, as suggested by our phenotypic clustering approach. Likewise, 3’UTR switching between paralogs/isoforms in other organisms will be necessary to validate our hypothesis for how specialized ribosomes are assembled.

## Materials and Methods

### Yeast strains, genomic manipulations, and growth conditions

Yeast strains, namely wild-type *S. cerevisiae* BY4741 and RP paralog (RPP) deletion mutants, were grown at 26°C, 30°C or 35°C in YPD (1% bacto–yeast extract, 2% bacto-peptone and 2% glucose), YPG (1% bacto–yeast extract, 2% bacto-peptone and 3% glycerol), YPO (1% bacto– yeast extract, 2% bacto-peptone and 0.2% oleic acid), or synthetic complete medium (SC), with or without selection. Conditional growth under stress was carried out using YPD media as control, 1.5M sorbitol for osmotic stress, 1.2M NaCl for high salt stress, and 5mM H_2_O_2_ for oxidative stress conditions.

### Preparation of the RP paralog knockout library

Most RPP knockout mutant strains were taken directly from the EUROSCARF deletion library (*RPP::KanMX*) (Supplementary Table 10; Source: EUROSCARF). The missing mutants were made by homologous recombination using standard PCR-based amplification of deletion knockout cassettes, using *HIS3* as a selection marker (Table 1; Source: This study). Sequences 40bp upstream of the start codon and 40bp downstream of the stop codon were selected as homology recombination regions for the knockout of RPP genes^47^. Deletion mutants were verified by colony PCR using a combination of specific forward primers for the gene (100-1000bp upstream of the start codon) and reverse primers against the downstream selection marker.

### C-terminus epitope tagging

Selected paralog pairs were FLAG-, HA-, or Myc-epitope-tagged at the carboxy terminus using a 40bp homologous region before and after the stop codon, followed by the epitope sequence with the stop codon and *URA3* gene as a selection marker ^47^. Subsequently, the *URA3* gene was removed by homologous recombination using 80 bp forward and reverse sequence oligos, including 40 bp sequences before and after the stop codon with the epitope and selection on 5-fluoroorotic acid (5-FOA) plates. Correct tagging was verified by PCR on genomic DNA.

### UTR switching

The length and sequences of 3’UTRs were obtained from the Saccharomyces Genome Database (SGD, https://www.yeastgenome.org/) First, the 3’UTR sequence of the RP paralogs was separated by inserting the *URA3* gene just after the stop codon. Next, the inserted *URA3* gene and 3’UTR of the paralog were removed, inserting the other paralog 3’UTR sequence with a homologous sequence of 40 bp length on both sides using homologous recombination and selected on 5-FOA plates. Correct UTR switching was verified by PCR on genomic DNA.

### Growth tests on liquid media

A Freedom EVO platform (Tecan Diagnostics, Switzerland) was used to automate the culturing and growth of yeast strains cultured in 96-well plates. The automation program included starter culture, inoculum dilution in fresh plate, and continuous culture to monitor growth. The starter culture of yeast strains was grown at 30°C with shaking at 1200 rpm for 20 sec at 40 min intervals for 30 h. Following this, 15µl of the starter culture was added to 135µl of fresh YPD medium (Control) with or without 1.5M D-Sorbitol (Osmotic Stress), 1.2M NaCl (High Salt Stress) or 4mM H_2_O_2_ (Oxidative Stress) and incubated at 30°C (A longer growth time at 26°C and a faster evaporation rate at 35°C limit this experiment to 30°C only). Plates were shaken on a shaker (BioShake) set at 1200 rpm for 20 sec prior to each O.D._600_ measurement at 40 min intervals using Tecan Infinitive plate reader. O.D._600_ measurements were pooled into one dataset, the growth curve was analyzed, and hierarchy clusters were plotted using tidyverse, growthcurver, and ggplot2 packages, respectively, in R^48–50^.

### Growth tests on solid media

For the drop test, 45 µl of overnight cultures (OD_600_ = 0.6-0.8) in 96-well plates were transferred using a liquid handler robot (LiHa) (Tecan) to 384 well plates. The three adjacent wells were subjected to serial dilutions of 1:5 for a total of four concentrations for each deletion strain. Cells were seeded on YPD, YPG, or YPO agar plates using a Singer pinning robot (Singer Instruments and Control). The plates were incubated at 26, 30, and 35°C and photo-documented after 2-4 days, 3–5 days, or 4-6 days, respectively. The colony area was calculated using an in-house ImageJ script as follows: images were first aligned, the background removed, and the colony area calculated using ImageJ’s analyze particles option. Finally, the measurement files were filtered, analyzed, and plotted in R using tidyverse and ggplot2 packages.

### PUNCH-P technique

Polysome isolation was performed as previously described ^13,51^. Briefly, yeast was grown in 500ml YPD or YPO at 30°C until the mid-log phase (OD_600_ = 0.6–0.8). Cells were washed once with ice-cold double distilled water, snap-frozen in liquid nitrogen, and stored at −80°C. Frozen yeast cells were resuspended in PEB and lysed using glass beads (0.4-0.6 mm). The lysates were centrifuged at 17,400 x *g* for 30 min at 4°C to remove cell debris. Subsequently, 6 ml of supernatant were loaded onto a 2.5ml 70% sucrose cushion. Samples were centrifuged at 60,000 rpm at 4°C for 2 h on an SW65 Ti ultracentrifuge rotor (Beckman Coulter, California, USA). The pellets were gently washed by dispensing and removing 500 μl of ice-cold RNase-free water. Ribosomes were suspended in 90µl PEB (without detergent) snap-frozen and stored at −80°C. The nascent polypeptide chains were purified by Biotin-dc-Puromycin labeling and pooling using streptavidin beads as previously described ^38^. For each sample, 20 OD_254_ units of polysomes were incubated with 100 pmol of biotin-dc-puromycin (Jena Bioscience, Germany) per OD_254_ unit at 37°C for 15 min. For each sample, 5µl of streptavidin beads (GE Healthcare, Chicago, USA) per OD_254_ were added and supplemented with 1 ml of SDS urea buffer (50 mM Tris-HCl, pH 7.5, 8M urea, 2% [wt / vol] SDS and 200mM NaCl) and incubated overnight at RT. The beads were washed with 1 ml of SDS - urea buffer and incubated in 1M NaCl for 30 min at 23°C, followed by five washes in Ultrapure water. The washed beads were subjected to mass spectrometric analysis (MS).

A total of 48 pulldowns of nascent polypeptides were analyzed by MS. Triplicate samples derived from cells (*e.g.* WT, *rpl12aΔ, rpl12bΔ, rpl19aΔ, rpl19bΔ, rpl34aΔ,* and *rpl34bΔ*) grown on either YPD or YPO media along with a negative control (NC; lacking biotin-dc-puromycin) were processed. Initial sample preparation involved tryptic digestion on beads aided by urea. Subsequent analysis employed nanoAcquity liquid chromatography (Waters Corp) coupled with Q Exactive HF MS (Thermo Fisher Scientific), evaluating each sample independently in a random order. Data processing for each of sample group was performed separately with the raw data processed using the MaxQuant v1.6.0.16 platform and analysis packages ^52^. Samples were screened against the *Saccharomyces cerevisiae* proteome using the Andromeda search engine and quantitative assessments facilitated using Perseus v1.6.0.7, which retained proteins with >2-fold, in at least one samples, then in the NC (to remove non-specific bead-binding proteins). Missing values were addressed using a low-value normal distribution. Hierarchical clustering and PCA were applied across paralog pairs, WT samples, and inter-pair assessments, aiding outlier identification and removal, with the two most reliable replicates selected for differential translatome analysis. Transformation of log values to their original form was undertaken, serving as input for DEP^53^, an R package designed for MaxQuant output of MS data. DEP enabled differential translatome analysis, accompanied by visualization tools such as volcano plots and hierarchical clustering. This comprehensive methodology underpins the robustness and interpretability of the ensuing results.

### Ribosome sucrose gradient fractionation

Sucrose gradient preparation: different concentrations of sucrose were prepared by adding 50%, 45%, 40%, 35%, 30%, 25%, 20% and 15% sucrose to polysome extraction buffer (PEB; 20 mM Tris-HCl, pH 7.4, 140 mM KCl, 10 mM MgCl2, 0.5 mM DTT, 2 µg/ml leupeptin, 40 U/ml RNAsin, 1.4 µg/ml pepstatin, 0.2 mg/ml heparin [Sigma-Aldrich], EDTA-free complete protease inhibitor mix [Roche] and 1% Triton X-100). Using the robotic liquid handler (LiHa; Tecan, Switzerland), 1.6 ml of each fraction was very slowly layered in a 14 ml ultracentrifuge tube (Beckman Coulter, California, USA), starting from higher to lower concentrations. Then the tubes were kept at 4°C for overnight to two days for diffusion. The diffused sucrose gradients were used for ribosome fractionation.

Yeast strains were grown overnight to saturation in YPD. 0.2 OD_600_ units of yeast cells were inoculated in 100-250 ml of YPD or YPO and cultured until the culture reached mid-log phase (OD_600_ = 0.6–0.8) at 30°C with constant shaking. Cells were incubated with cycloheximide for 5 min just before harvesting. The yeast cells were washed once with double distilled ice-cold water and snap frozen in liquid nitrogen. Samples were stored at −80°C for future use or suspended in PEB buffer for immediate use. Cells were lysed with glass beads at 4°C for 10 min, and cell debris was removed by centrifugation at 15,000xg, 4°C. The supernatant was layered on top of the 50-15% linear sucrose gradient. The samples were ultracentrifuged at 39,000 rpm at 4°C for 2.5 h in an SW41 rotor (Beckman Coulter, California, USA). Finally, 135ul of fractions per well were collected on a 96-well UV transparent plate from the top using 200ul LiHa. The OD260 was measured for each well using the Tecan plate reader. Fractions corresponding to the small subunit (40S), large subunit (60S), monosomes (80S), and polysomes were pooled for downstream experiments.

### 80S monosome composition analysis

250 ml of mid-log C-terminal FLAG-tagged and untagged yeast cells were lysed. The 80S monosome fraction was collected from the sucrose gradient, as mentioned above in Ribosome sucrose gradient fractionation method. The collected monosomes were then 1:1 diluted with PEB and centrifuged at 60,000 rpm at 4°C overnight on an SW65 Ti ultracentrifuge rotor (Beckman Coulter, California, USA). The pellets were resuspended in 250µl PEB with 50 mM NH_4_Cl. The concentration of monosomes was measured at OD_260_. An equal amount of monosomes per sample was incubated overnight with 100 µl of anti-FLAG Sepharose beads (Cell Signaling Technology, Massachusetts, USA) at 4°C. The samples were pelleted at 1000xg and washed five times with PEB with 50 mM NH_4_Cl. Finally, samples were eluted with at least 50 ul of FLAG peptide (1mg/ml) in PEB buffer with 50 mM NH_4_Cl and sent for MS or used for Western blotting.

Monosome composition analysis was conducted using a methodology like that used for PUNCH-P, however, we employed in-solution digestion instead of on-bead digestion. Data processing was performed separately for each set of growth media samples belonging to the different FLAG-tagged RP paralogs. As done for the PUNCH-P MS, we conducted quantitative analysis to assess differential composition between ribosomal paralogs. A scatter plot was generated using the ggplot2 R package to visualize differences in composition between paralog pairs. Additionally, we combined the YPO and YPD datasets to explore qualitative variations in composition. For this purpose, we employed the Pheatmap and Dendextend R packages to conduct Hierarchical clustering and Tanglegram analysis, respectively ^32^.

### Peroxisome induction

Yeast cells expressing the peroxisomal marker PTS-RFP were grown overnight in YPD. 0.2 OD_600_ cells were inoculated in peroxisome induction medium (0.3% yeast extract, 0.5% peptone, 0.5% KH_2_PO_4_, pH 6), containing 0.225% oleate, and grown overnight at 30°C. The cells were then subjected to further experiments.

### Image stream analysis and confocal microscopy

Following peroxisome induction, cells were washed once with TE buffer and immediately taken for flow cytometric analysis or photo documentation ^54^. The fluorescence signal readout was recorded using an ImageStreamX imaging flow cytometer (Amnis, Seattle, USA) and analyzed using IDEAS software (Merck Millipore). For confocal microscopy, images were acquired at 26°C using an LSM800 confocal microscope equipped with a Plan Apochromat 100×1.40 NA oil objective (Zeiss, Germany). Images were analyzed using ImageJ (NIH, USA).

### Western Blotting

Total cell lysates, ribosome fractions, or immunoprecipitants were separated by SDS-PAGE. The proteins were transferred to a 0.45 μm pore size cellulose membrane (10401180, GE Healthcare), then probed with anti-HA (901514, 1:5000, BioLegend, San Diego, CA); anti-FLAG (F3165, 1:5000, Sigma); or anti-Myc (9E10, 1:200, Santa Cruz) antibody. The tagged proteins were detected using HRP conjugated sheep anti-mouse or anti-rabbit IgG antibody (1:10000, GE Healthcare, Uppsala, Sweden) and visualized using the Amersham ECL Western blot detection reagent (GE Healthcare), according to the manufacturer’s protocol in an ImageQuant LAS 4000 mini (GE Healthcare). Finally, the band intensity was measured using ImageStudio, and graphs were plotted in R.

### Polysome proteomics

Formaldehyde crosslinking: Yeast strains were grown overnight to saturation in YPD and 0.1 OD600 units of cells were inoculated in 50 mL of YPD or YPO. Cultures were grown to mid-log phase (OD600 ∼0.8) at 30°C with constant shaking. Formaldehyde crosslinking was carried out as in Crawford *et al*., 2022^45^. Briefly, cultures were crosslinked with 0.8% formaldehyde for 1 h on ice. Excess formaldehyde was quenched with glycine and cells were harvested by centrifugation and lysed by vortexing with glass beads into 200 µl of polysome lysis buffer (20 mM HEPES pH 7.4, 2 mM magnesium acetate, 100 mM potassium acetate, 0.5 mM DTT, 0.1% DEPC).

Polysome profiling & protein extraction: Polysome profiling and protein extraction were carried out as in Crawford et al, 2022. Briefly, 1.5 A260 units of lysate were layered on to 15-50% sucrose gradients prepared in 12 ml thin-walled open polyallomer tubes (Seton Scientific) and separated by ultracentrifugation in an SW41 Ti rotor (2.5 hrs at 278,000 x *g*, 4°C). Fractions corresponding to monosomes, ‘light polysomes’ (2-4 ribosomes per mRNA) and ‘heavy polysomes’ (≥5 ribosomes per mRNA) were collected manually for mass spectrometry analysis. Protein was precipitated from the fractions by adding a half volume of 40% trichloroacetic acid (TCA) and incubating overnight. Protein pellets were washed with acetone and air-dried.

Trypsin digestion: Pellets were resuspended in 25 mM ammonium bicarbonate (AmBic) to a volume of 80 µl. ‘Total’ samples from the original cell lysates (corresponding to 0.15 A260 units) were made up to a volume of 80 µl using 25 mM AmBic. RapiGest SF Surfactant (Waters; 1% w/v in 25 mM AmBic, final concentration 0.012%) was added and samples were incubated for 10 min at 80°C, with shaking at 450 rpm. Samples were reduced by adding DTT (72 mM in 25 mM AmBic, final concentration 4 mM) and incubating for 10 min at 60°C, with shaking at 450 rpm. Samples were cooled to room temperature then alkylated by adding iodoacetamide (266 mM in 25 mM AmBic, final concentration 14 mM) and incubating for 30 min at room temperature in the dark. Excess iodoacetamide was quenched by adding further DTT (final concentration 7 mM, including the first addition) and vortexing. Samples were digested by adding Sequencing Grade Modified Trypsin (Promega; 400 µg/ml in 25 mM AmBic, final concentration 19 µg/ml) at an enzyme-to-protein ratio of 1:50 and incubating for 16 h at 37°C. Samples were acidified by adding trifluoroacetic acid (TFA; final concentration 0.5% v/v) and incubating for 45 min at 37°C. Samples were cleared by centrifuging for 15 min at 13,000 x *g*, 4°C and the supernatant was transferred to a fresh tube for desalting. OLIGO R3 Reversed-Phase Resin (Thermo Scientific) in a FiltrEX 96-well Filter Plate with 0.2 µm PVDF Membrane (Corning) was used for desalting. Peptides were bound to the resin for 5 min at room temperature, with shaking at 500 rpm, and the supernatant was removed by centrifugation for 1 min at 200 x g. Peptides were washed twice with 0.1% aqueous formic acid, eluted twice by adding 0.1% formic acid in 30% acetonitrile, then were dried to completeness in a SpeedVac SPD1010 vacuum centrifuge (Thermo Scientific).

Mass spectrometry: Liquid chromatography (LC) was carried out using an UltiMate 3000 Rapid Separation Binary System (Thermo Fisher Scientific). Peptides were concentrated using an ACQUITY UPLC M-Class Symmetry C18 Trap Column (180 μm inner diameter, 20 mm length (Waters)) and then separated using an ACQUITY UPLC M-Class Peptide BEH C18 Column (75 μm inner diameter, 250 mm length, 1.7 μm particle size (Waters)). A gradient starting with 99% Buffer A (0.1% formic acid in water) and 1% Buffer B (0.1% formic acid in acetonitrile) and increasing to 75% Buffer A and 25% Buffer B was used to separate the peptides over 45 min at a flow rate of 200 nL/min. Label-free tandem MS was performed using a Q Exactive HF mass spectrometer (Thermo Scientific). Peptides were selected for fragmentation and MS2 analysis automatically by data-dependent analysis.

MS data analysis: Raw MS data were processed using MaxQuant version 2.0.3.0 ^52^. A peptide mass tolerance of 20 ppm was used for the first search, 4.5 ppm for the main search, and 0.5 Da for the MS/MS fragment ions. The peak list was searched against the Uniprot *Saccharomyces cerevisiae* database (accessed 10th February 2017) using the built-in Andromeda search engine^55^. Peptide-spectrum matches and protein groups were each filtered at a false-discovery rate of 1%. Data analysis was performed using R (version 4.2.2)^56^ and the packages MSstats (version 3.18.5)^57^, protti (version 0.6.0)^58^, Mfuzz (version 2.56.0)^59^ and clusterProfiler (version 4.4.4)^60^. Plots were made using ggplot2 (version 3.4.3)^48^.

## Data availability

Translatome, monosome, polysome, and whole cell proteomics data have been deposited to the ProteomeXchange consortium via the PRIDE partner repository^61^ with the following dataset identifiers and access information: Yeast polysome composition during glucose and oleate growth (accession #PXD046823; username, password: m5ZmCoWd); Translatome profiling of WT, rpl12bΔ, and RPL12A-B3’UTR cells grown on oleate-containing medium (accession #PXD050699; username; password: 2XniEa7f); Translatome profiling of WT, rpl12bΔ, and RPL12A-B3’UTR cells grown on oleate-containing medium (accession #PXD050699; username, password: 2XniEa7f); Translatome profiling of yeast during glucose and oleate growth (accession #PXD050697; username, password: nNcHwLEM); Yeast ribosomal protein paralog-specific monosome composition during glucose and oleate growth (accession #PXD050695; username, password: THdgoJuy)

## Materials availability

Yeast strains used in this study (Supplementary Table 10) are available upon request.

## Acknowledgements

The authors thank Drs. Amir Prior and Yishai Levin of the Mass Spectrometry Unit of the INCPM (Weizmann Institute of Science) for MS experiments and analysis. We thank David Knight and colleagues at the University of Manchester BioMS Core Facility (RRID:SCR_020987) for processing mass spectrometry samples and their helpful advice. The authors also thank Dr. Rohini R. Nair for pRS316-Rpl19/34a/b plasmids, Prof. Maya Schuldiner for Pex25-BirA/Rpl16-HA-AVI strain, Dr. Tsviya Olender for bioinformatic help, and Dr. Gal Haimovich for a critical reading of the manuscript. This work was funded by grants to J.E.G. from the Israel Science Foundation (#980/22) and to J.E.G. and G.D.P. from the Weizmann-UK Joint Research Program. J.E.G. holds the Besen-Brender Chair in Microbiology and Parasitology, Weizmann Institute. R.S. is presently located at the New York Genome Center.

## Author Contributions

J.E.G. conceptualized the study. R.S., R.C. G.P., and J.E.G. conceived the experiments and wrote the manuscript. R.S., R.C., and D.P.M. performed the experiments, related data analyses, and data curation. J.E.G. and G.P. secured funding.

## Supplementary Figure Legends

**Supplementary Figure 1.**
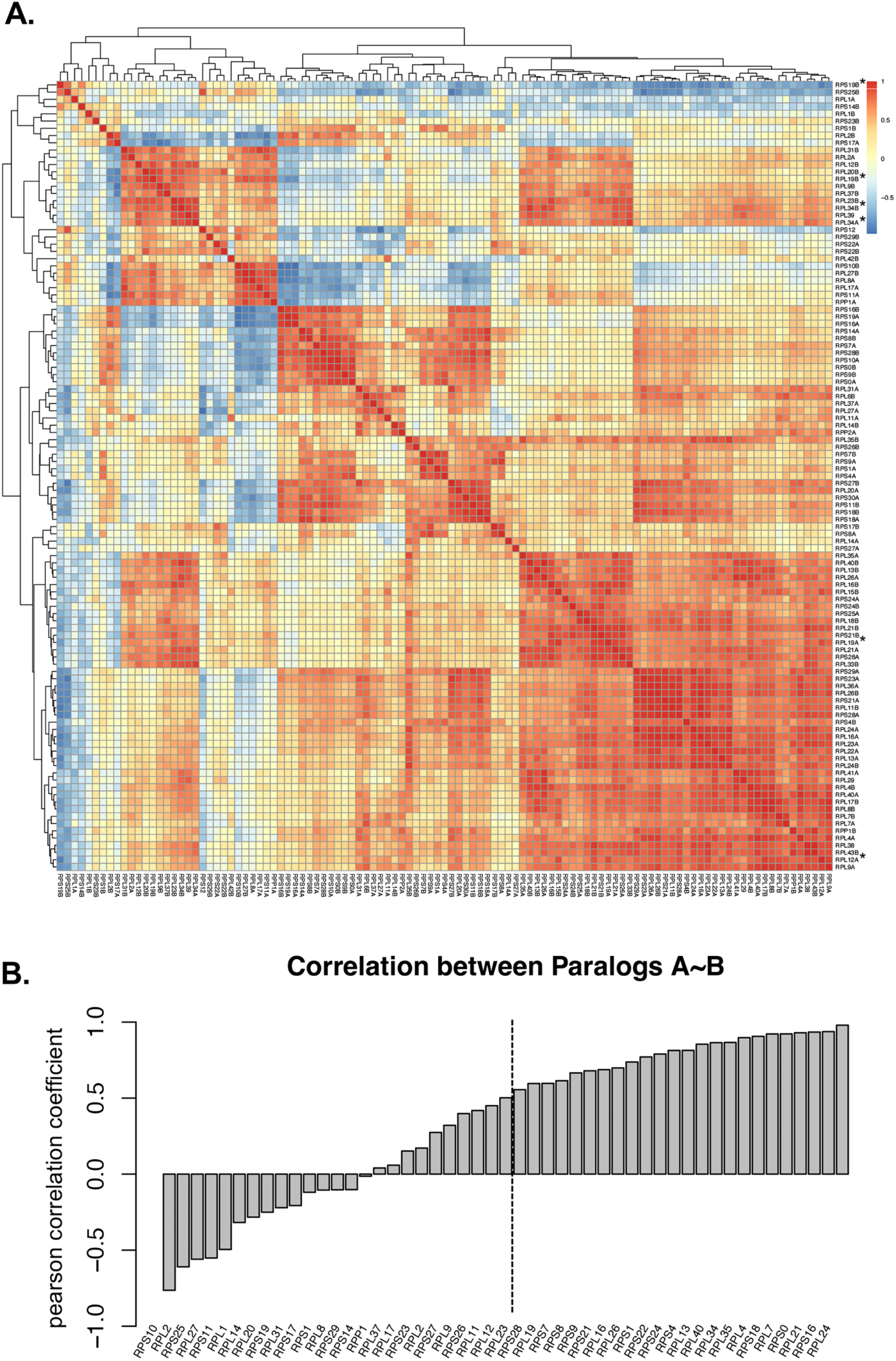
Correlative analysis of the growth phenotype of the RP deletion strains on glucose-, glycerol-, and oleate-containing media (A) Heatmap of the correlation matrix for all strains of the RP gene deletion library grown on the different carbon sources (glucose, glycerol, and oleate). Note that the RP paralog deletion strains form distinct phenotypic clusters. (B) Correlation between paralogs of the same ribosomal protein. Note that 28 out 54 the paralogs show either anticorrelation or a lower correlation (*p* correlation coefficient <0.5) under the different carbon source growth conditions.

**Supplementary Figure 2.**
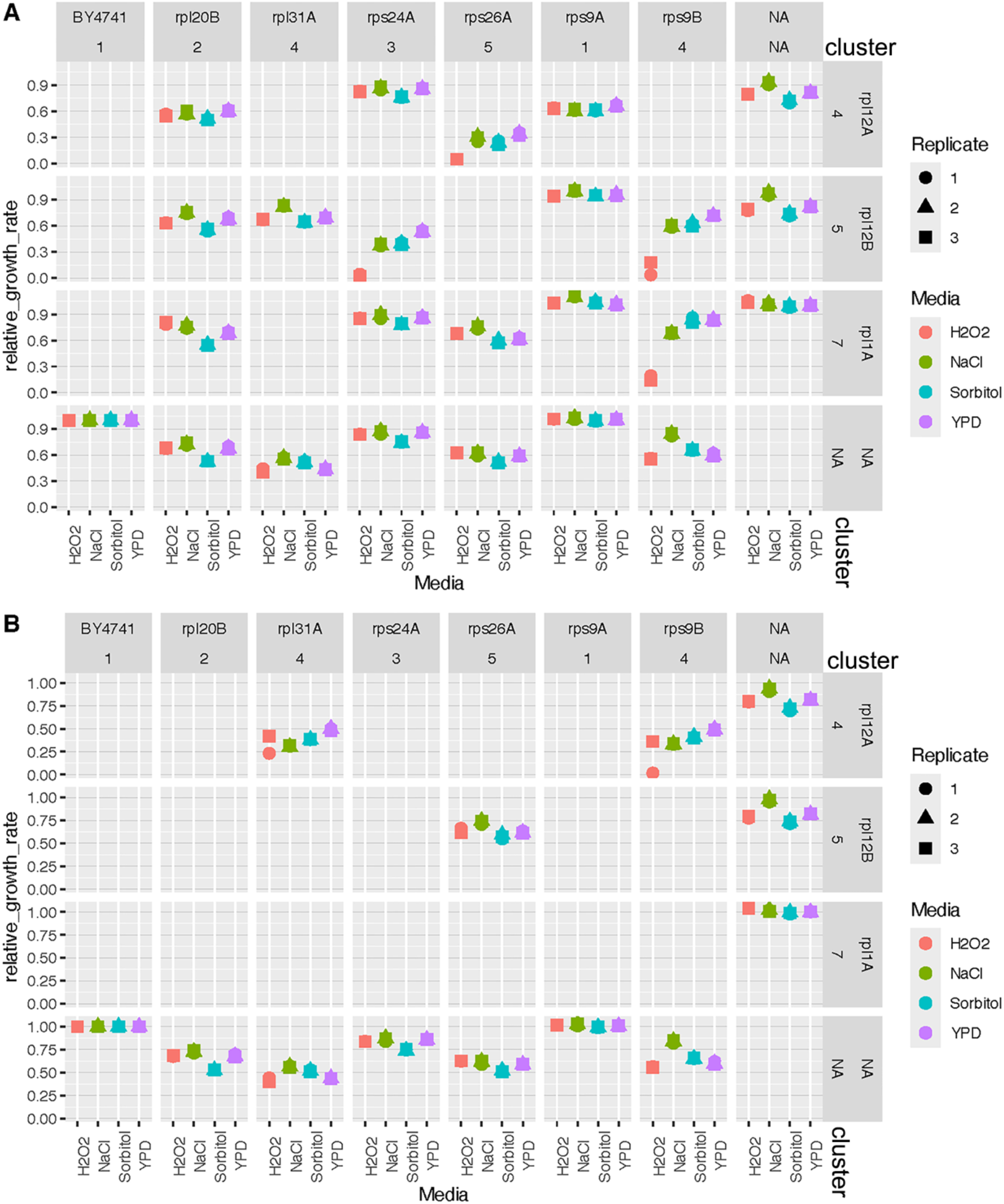
Synthetic interaction of phenotypic clusters (A) Graph of the relative growth rates (r) of single RP paralog deletions (NA) or pair-wise intercluster RP double deletion mutants, as compared to BY4741 WT cells, under different stress conditions (YPD - normal, 1.2M NaCl - high salt, 1.5M Sorbitol - osmotic, and 5mM H_2_O_2_ – oxidative stress). Results from three biological replicas are shown. (B) Same as in *A*, but showing relative growth rates of single RP paralog deletions (NA) or pair-wise intracluster RP double deletion mutants, as compared to BY4741 WT cells.

**Supplementary Figure 3.**
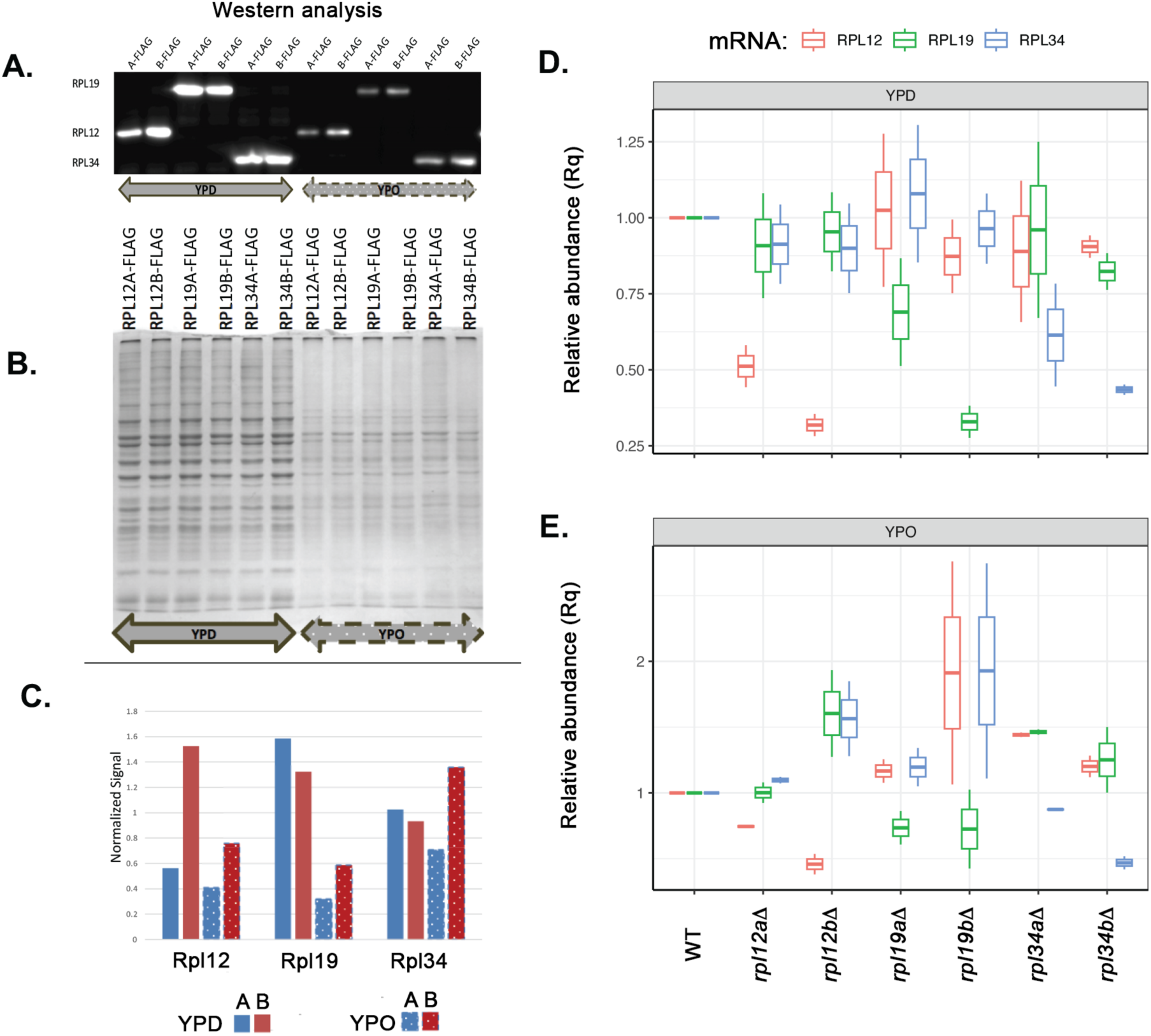
RP paralog protein expression on glucose- and oleate-containing media (A) Immunoblot of total cell lysates of the C-terminal FLAG-tagged RP paralogs in the WT background grown on glucose-containing (YPD) or oleate-containing medium (YPO) and probed with anti-FLAG antibody. (B) Coomassie staining of SDS-PAGE gel used for the immunoblot shown in *A*. (C) Quantification of the Western blot shown in *A* by densiometric analysis, as normalized with total protein. A and B indicate the FLAG-tagged paralog expressed. (D,E) Quantification of the relative abundance of RP mRNAs in *rpl12aΔ, rpl19aΔ, rpl34aΔ, rpl12bΔ, rpl19bΔ*, and *rpl34bΔ* cells relative to WT cells, as quantified by RT-qPCR analysis. mRNA abundance was normalized to the *ACT1* gene and the cells were grown till mid-log phase on YPD (D) and YPO medium (E), respectively.

**Supplementary Figure 4.**
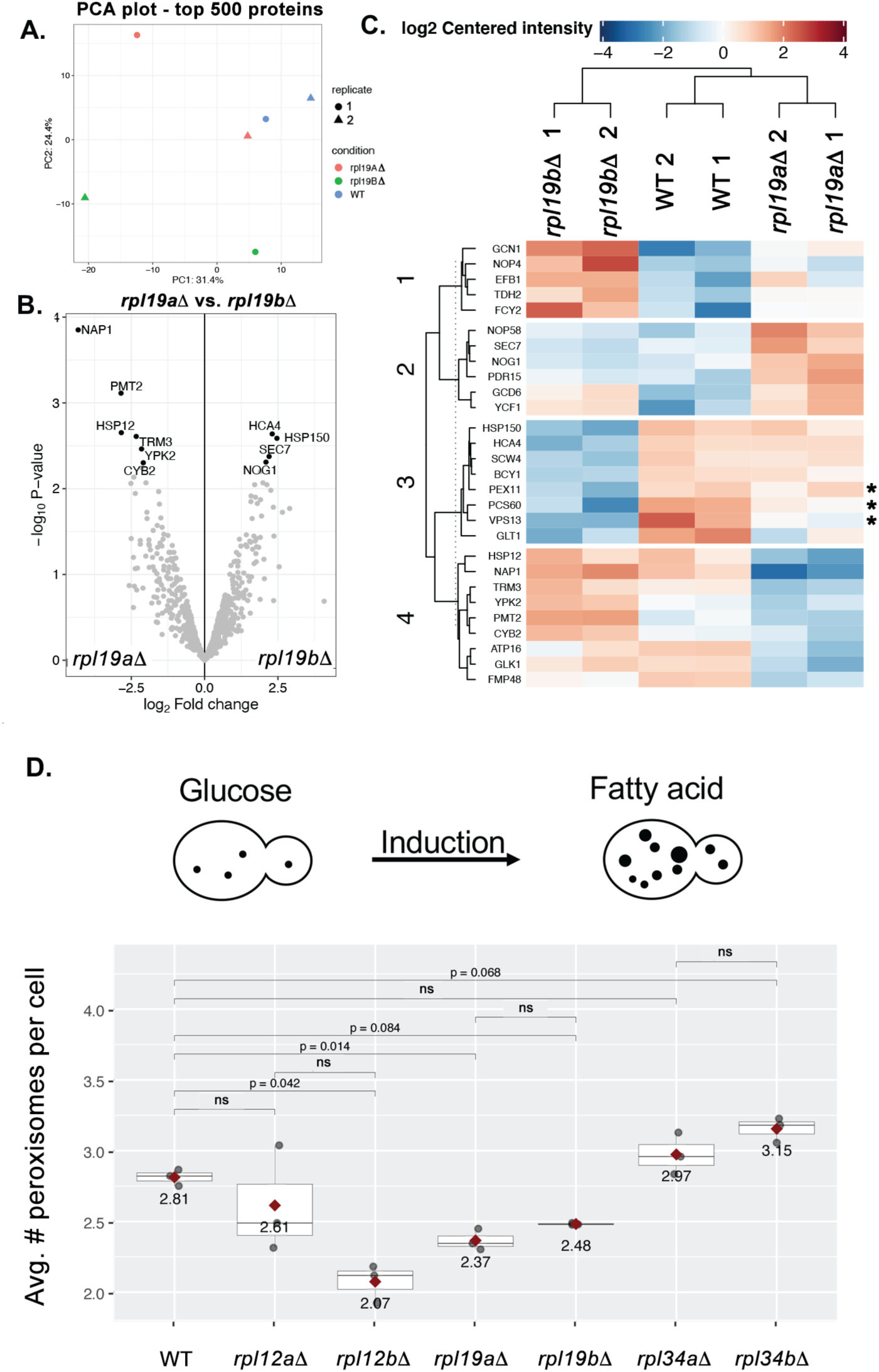
rpl19ϕ translatome profiling and peroxisome induction of the RP deletion strain grown on oleate-containing medium. (A) Principal component analysis of the translatomes (top 500 proteins) of WT, *rpl19aΔ,* and *rpl19bΔ* cells grown on oleate-containing medium (YPO). Results of two biological replicas are shown. (B) Volcano plot of the PUNCH-P translatomes derived from WT, *rpl19aΔ,* and *rpl19bΔ* polysomes from cells grown on YPO (C) Heat map of clustering of differentially-translated nascent polypeptides derived from WT, *rpl19aΔ,* and *rpl19bΔ* polysomes from cells grown on YPO. Asterisks indicate peroxisomal and peroxisome-related proteins. (D) Quantification of the number of peroxisomes per cell in WT and the different paralog deletion strains, as scored by counting GFP-SKL labeled peroxisomes using fluorescence microscopy. Peroxisomes were scored after 12hrs of induction on YPO media.

**Supplementary Figure 5.**
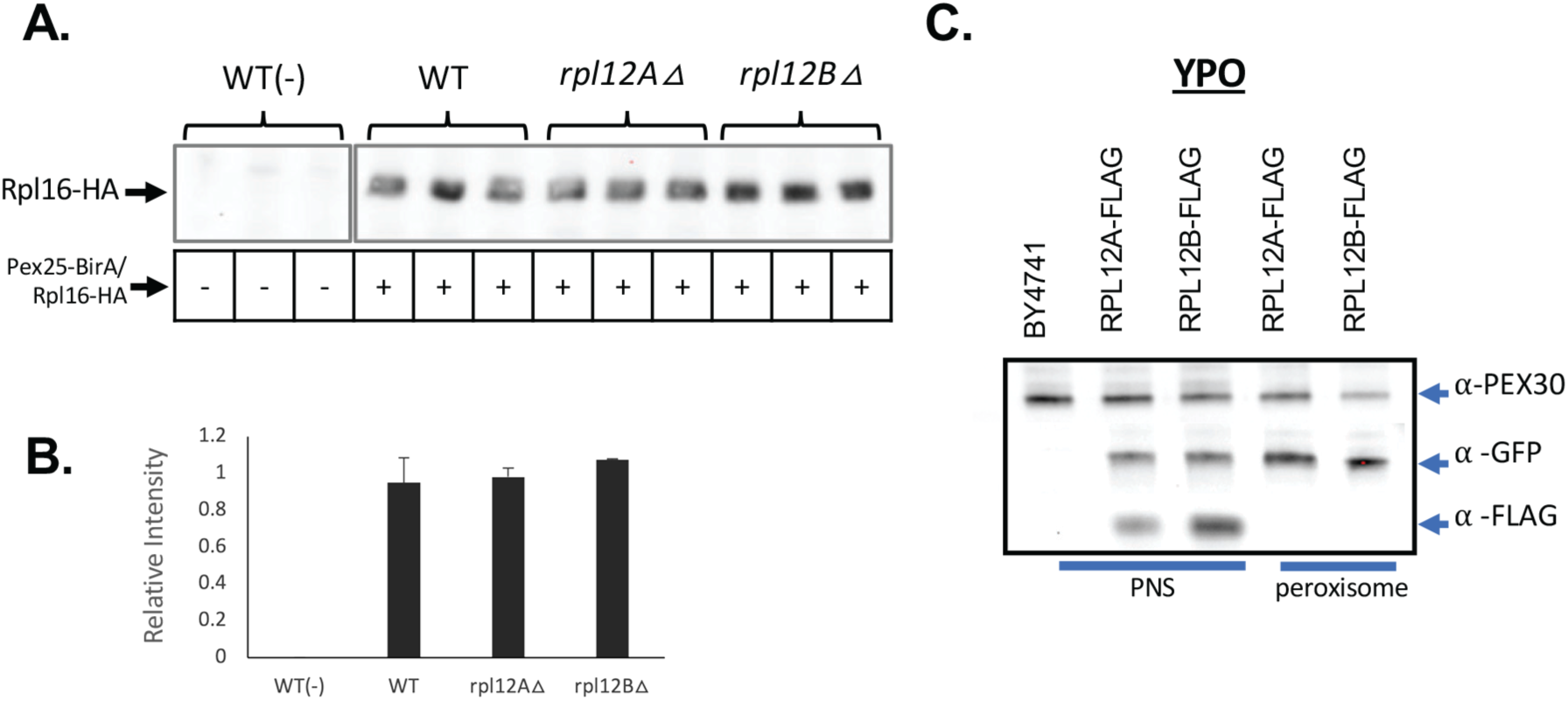
Examination of paralog-specific localized translation associated with peroxisomes (A) Proximity labeling assay using BioID. WT, *rpl12aΔ,* and *rpl12bΔ* cells expressing a peroxisomal membrane protein (Pex25) fused to BirA and both Rpl16a and Rpl16b fused to the HA and AviTag epitopes were used for BioID labeling. Unlabeled WT cells were used as a control. Cells were grown on oleate-containing medium (YPO). Shown is Western blot analysis of total cell lysate of WT, *rpl12aΔ,* and *rpl12bΔ* cells probed with streptavidin-HRP to quantify the association of Rpl16-HA-containing ribosomes proximal to the peroxisome. (B) Quantification of the results shown in *A.* Band intensity for each signal was divided by the mean intensity measured for all samples to show variation. (C) Co-immunoprecipitation and Western blot analysis of the peroxisome fraction. Immunoblot of post-nuclear supernatant (PNS) and peroxisome fraction (peroxisome) derived from WT BY4741 cells, Rpl12A-FLAG cells expressing GFP-SKL, and Rpl12B-FLAG cells expressing GFP-SKL grown on YPO medium. Cells were grown on YPO medium prior to processing and subsequent sucrose- and nycodenz density gradient centrifugation of the PNS fraction to obtain the peroxisome fraction. Immunoblots were probed with anti-Pex30 (top), anti-GFP, and anti-FLAG antibodies. Note that we did not observe ribosome association with the peroxisome fraction, as assessed using Rpl12a-FLAG or Rpl12b-FLAG.

**Supplementary Figure 6.**
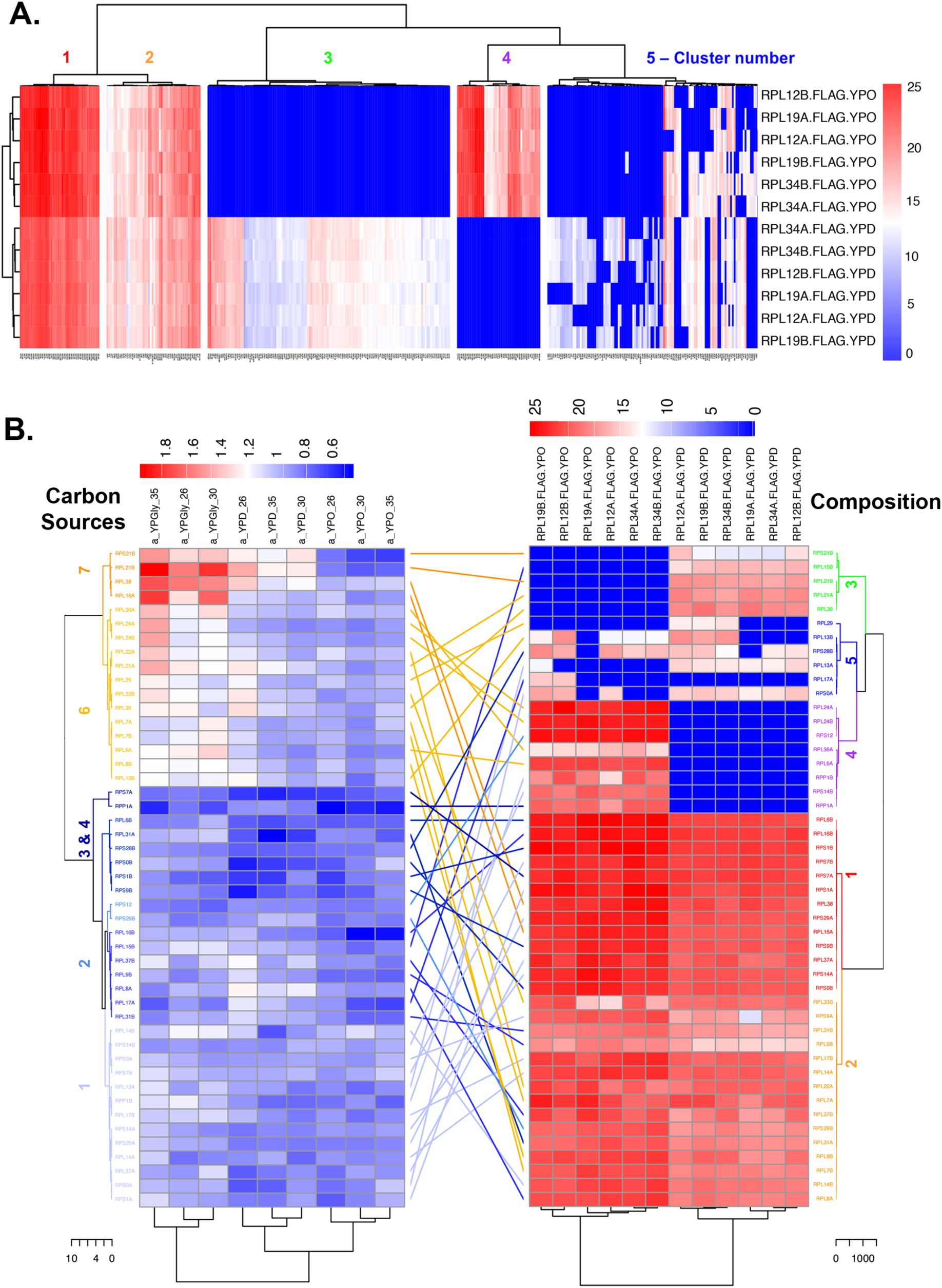
Comparative analysis of protein composition versus phenotype in functionally distinct ribosomes (A) Hierarchical clustering of the protein composition of RP paralog-specific monosomes derived from cells grown in YPD and YPO at 30°C. (B) Tanglegram of the heatmaps for hierarchal clustering of the growth phenotype on the different carbon sources (left) and that of the monosome protein composition (right). The dendrograms are color-coordinated based on cluster number, labeled according to the carbon source clusters (see Figure 2A as well).

**Supplementary Figure 7.**
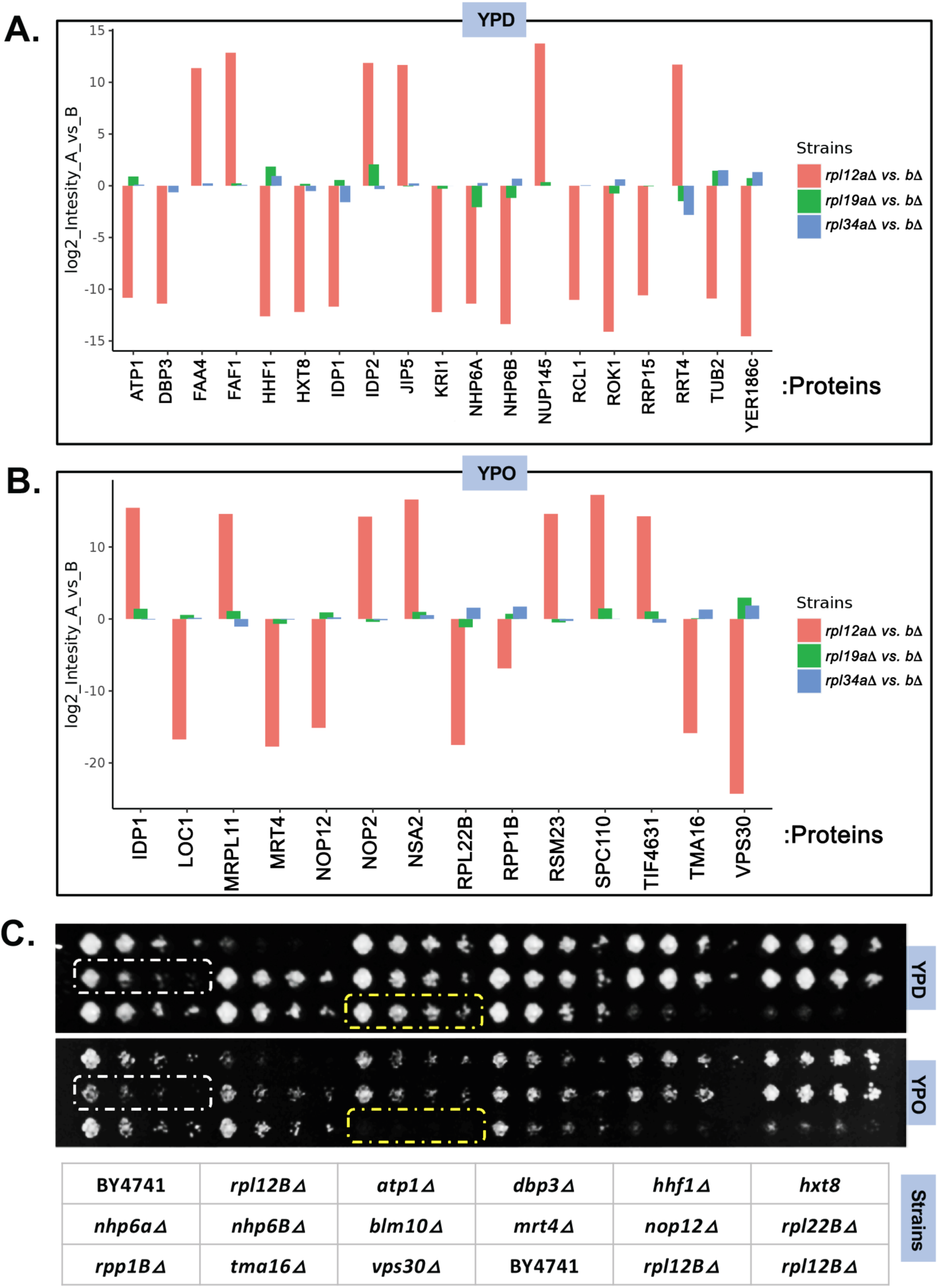
RAP abundance in paralog-specific ribosomes and the phenotype of their deletion in WT cells (A and B) Bar graph of the levels of RPs and RAPs that were differentially associated with the Rpl12a-FLAG and Rpl12b-FLAG ribosomes (Figure 5D), as measured in the *rpl12ϕ, rpl19ϕ,* and *rpl134Δ a* and *b* paralog deletion strains grown either on YPD (A) or YPO (B). (C) Drop test of the deletions of genes encoding RPs and RAPs that interacted specifically with the Rpl12b-FLAG ribosome in Figure 5D. Gene deletions were made in WT cells, which were grown to mid-log phase in YPD (top) or YPO (middle), prior to serial dilution and plating by drops onto solid medium. Bottom matrix shows the name and location of WT cells and each deletion strain. The deletion of *VPS30*, which encodes the most enriched RAP in Rpl12b-FLAG ribosomes (B) does not grow on YPO (C), unlike on YPD.

**Supplementary Figure 8.**
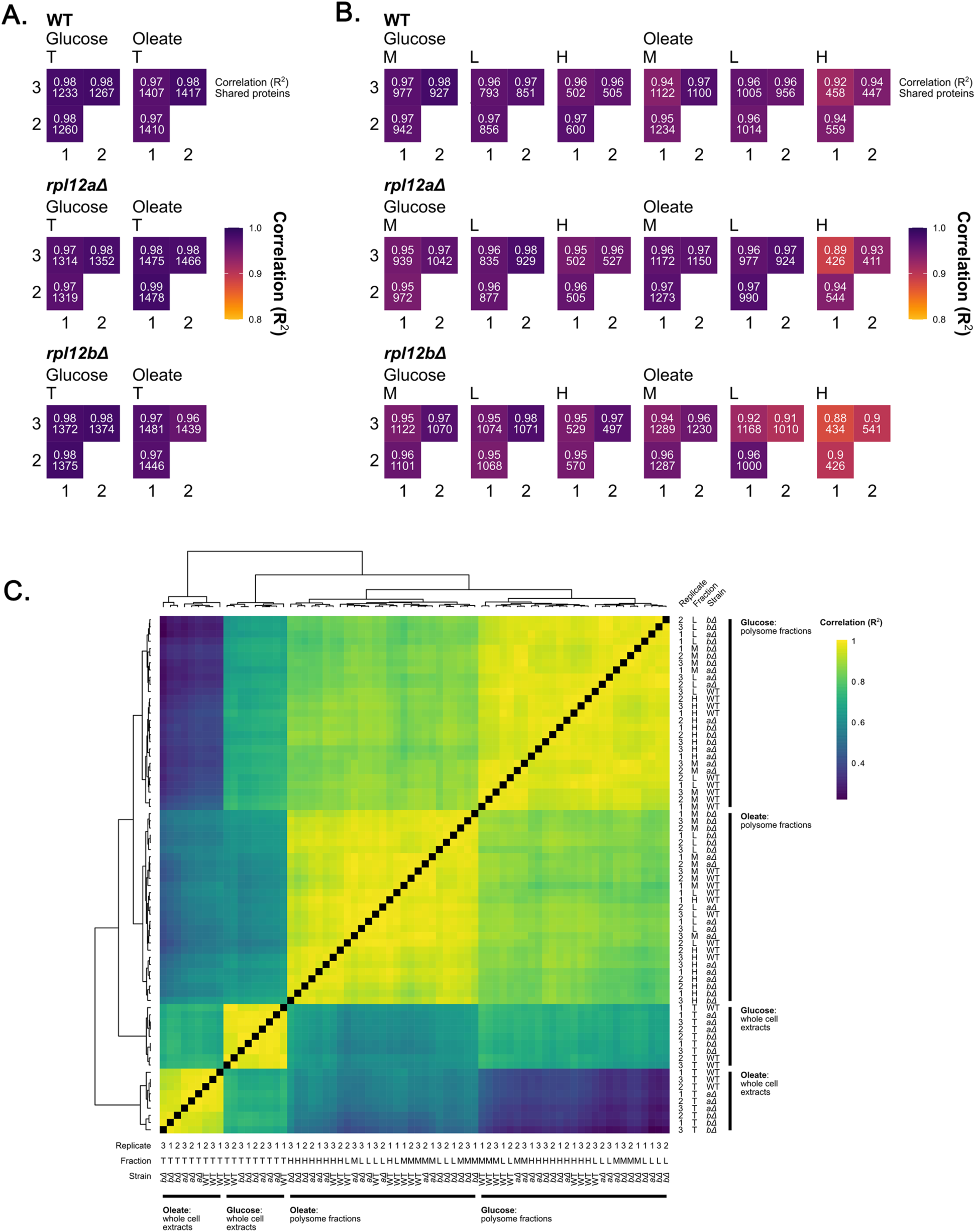
Different growth media induce changes in the protein composition of polysome fractions and the total proteome (A and B) Comparison of proteomic data replicates for (A) whole cell extracts and (B) polysome fractions. The exact R^2^ value and the number of shared proteins used for linear regression are shown for each comparison. (C) Comparison of proteomic data between all replicates of all samples. Samples were ordered by hierarchical clustering, shown in the dendrograms. *aΔ* – *rpl12aΔ*, *bΔ* – *rpl12bΔ*. T – total, M – monosome, L – light polysomes, H – heavy polysomes.

## References

1. Gay, D.M., Lund, A.H., and Jansson, M.D. (2022). Translational control through ribosome heterogeneity and functional specialization. Trends Biochem Sci 47, 66–81.

2. Genuth, N.R., and Barna, M. (2018). Heterogeneity and specialized functions of translation machinery: from genes to organisms. Nat Rev Genet 19, 431–452.

3. Gerst, J.E. (2018). Pimp My Ribosome: Ribosomal Protein Paralogs Specify Translational Control. Trends Genet 34, 832–845.

4. Guo, H. (2018). Specialized ribosomes and the control of translation. Biochem Soc Trans 46, 855–869.

5. Perbal, B. (1980). Transformation phenotype of polyoma virus-transformed rat fibroblasts: plasminogen activator production is modulated by the growth state of the cells and regulated by the expression of an early viral gene function. J Virol 35, 420–427.

6. Li, D., and Wang, J. (2020). Ribosome heterogeneity in stem cells and development. J Cell Biol 219.

7. Dias-Fields, L., and Adamala, K.P. (2022). Engineering Ribosomes to Alleviate Abiotic Stress in Plants: A Perspective. Plants (Basel) 11, 2097.

8. Ferretti, M.B., Ghalei, H., Ward, E.A., Potts, E.L., and Karbstein, K. (2017). Rps26 directs mRNA-specific translation by recognition of Kozak sequence elements. Nat Struct Mol Biol 24, 700–707.

9. Ghulam, M.M., Catala, M., and Abou Elela, S. (2020). Differential expression of duplicated ribosomal protein genes modifies ribosome composition in response to stress. Nucleic Acids Res 48, 1954–1968.

10. Imami, K., Milek, M., Bogdanow, B., Yasuda, T., Kastelic, N., Zauber, H., Ishihama, Y., Landthaler, M., and Selbach, M. (2018). Phosphorylation of the Ribosomal Protein RPL12/uL11 Affects Translation during Mitosis. Mol Cell 72, 84–98.e9.

11. Jang, S., Lee, J., Mathews, J., Ruess, H., Williford, A.O., Rangan, P., Betrán, E., and Buszczak, M. (2021). The Drosophila ribosome protein S5 paralog RpS5b promotes germ cell and follicle cell differentiation during oogenesis. Development 148.

12. Kearse, M.G., Chen, A.S., and Ware, V.C. (2011). Expression of ribosomal protein L22e family members in Drosophila melanogaster: rpL22-like is differentially expressed and alternatively spliced. Nucleic Acids Res 39, 2701–2716.

13. Segev, N., and Gerst, J.E. (2018). Specialized ribosomes and specific ribosomal protein paralogs control translation of mitochondrial proteins. J Cell Biol 217, 117–126.

14. Shi, Z., Fujii, K., Kovary, K.M., Genuth, N.R., Röst, H.L., Teruel, M.N., and Barna, M. (2017). Heterogeneous Ribosomes Preferentially Translate Distinct Subpools of mRNAs Genome-wide. Mol Cell 67, 71–83.e7.

15. Farley, K.I., and Baserga, S.J. (2016). Probing the mechanisms underlying human diseases in making ribosomes. Biochem Soc Trans 44, 1035–1044.

16. Kapur, M., Monaghan, C.E., and Ackerman, S.L. (2017). Regulation of mRNA Translation in Neurons-A Matter of Life and Death. Neuron 96, 616–637.

17. Nakhoul, H., Ke, J., Zhou, X., Liao, W., Zeng, S.X., and Lu, H. (2014). Ribosomopathies: mechanisms of disease. Clin Med Insights Blood Disord 7, 7–16.

18. Narla, A., and Ebert, B.L. (2010). Ribosomopathies: human disorders of ribosome dysfunction. Blood 115, 3196–3205.

19. Stern-Ginossar, N., Thompson, S.R., Mathews, M.B., and Mohr, I. (2019). Translational Control in Virus-Infected Cells. Cold Spring Harb Perspect Biol 11, a033001.

20. Sulima, S.O., Kampen, K.R., and De Keersmaecker, K. (2019). Cancer Biogenesis in Ribosomopathies. Cells 8, 229.

21. Wang, X., Zhu, J., Zhang, D., and Liu, G. (2022). Ribosomal control in RNA virus-infected cells. Front Microbiol 13, 1026887.

22. Wong, Q.W.-L., Li, J., Ng, S.R., Lim, S.G., Yang, H., and Vardy, L.A. (2014). RPL39L is an example of a recently evolved ribosomal protein paralog that shows highly specific tissue expression patterns and is upregulated in ESCs and HCC tumors. RNA Biol 11, 33– 41.

23. Lapointe, C.P., Grosely, R., Johnson, A.G., Wang, J., Fernández, I.S., and Puglisi, J.D. (2021). Dynamic competition between SARS-CoV-2 NSP1 and mRNA on the human ribosome inhibits translation initiation. Proc Natl Acad Sci U S A 118.

24. Simeoni, M., Cavinato, T., Rodriguez, D., and Gatfield, D. (2021). I(nsp1)ecting SARS-CoV-2-ribosome interactions. Commun Biol 4, 715.

25. Tidu, A., Janvier, A., Schaeffer, L., Sosnowski, P., Kuhn, L., Hammann, P., Westhof, E., Eriani, G., and Martin, F. (2020). The viral protein NSP1 acts as a ribosome gatekeeper for shutting down host translation and fostering SARS-CoV-2 translation. RNA 27, 253–264.

26. McIntosh, K.B., and Warner, J.R. (2007). Yeast ribosomes: variety is the spice of life. Cell 131, 450–451.

27. Woolford, J.L., and Baserga, S.J. (2013). Ribosome biogenesis in the yeast Saccharomyces cerevisiae. Genetics 195, 643–681.

28. Yang, Y.-M., Jung, Y., Abegg, D., Adibekian, A., Carroll, K.S., and Karbstein, K. (2023). Chaperone-directed ribosome repair after oxidative damage. Mol Cell 83, 1527–1537.e5.

29. Mattaini, K.R., Brignole, E.J., Kini, M., Davidson, S.M., Fiske, B.P., Drennan, C.L., and Vander Heiden, M.G. (2015). An epitope tag alters phosphoglycerate dehydrogenase structure and impairs ability to support cell proliferation. Cancer Metab 3, 5.

30. Steffen, K.K., MacKay, V.L., Kerr, E.O., Tsuchiya, M., Hu, D., Fox, L.A., Dang, N., Johnston, E.D., Oakes, J.A., Tchao, B.N., et al. (2008). Yeast life span extension by depletion of 60s ribosomal subunits is mediated by Gcn4. Cell 133, 292–302.

31. Ohashi, Y. (2021). Class III phosphatidylinositol 3-kinase complex I subunit NRBF2/Atg38 - from cell and structural biology to health and disease. Autophagy 17, 3897–3907.

32. Galili, T. (2015). dendextend: an R package for visualizing, adjusting and comparing trees of hierarchical clustering. Bioinformatics 31, 3718–3720.

33. Dahan, N., Bykov, Y.S., Boydston, E.A., Fadel, A., Gazi, Z., Hochberg-Laufer, H., Martenson, J., Denic, V., Shav-Tal, Y., Weissman, J.S., et al. (2022). Peroxisome function relies on organelle-associated mRNA translation. Sci Adv 8, eabk2141.

34. Haimovich, G., Cohen-Zontag, O., and Gerst, J.E. (2016). A role for mRNA trafficking and localized translation in peroxisome biogenesis and function? Biochim Biophys Acta 1863, 911–921.

35. Zipor, G., Haim-Vilmovsky, L., Gelin-Licht, R., Gadir, N., Brocard, C., and Gerst, J.E. (2009). Localization of mRNAs coding for peroxisomal proteins in the yeast, Saccharomyces cerevisiae. Proc Natl Acad Sci U S A 106, 19848–19853.

36. Lockshon, D., Surface, L.E., Kerr, E.O., Kaeberlein, M., and Kennedy, B.K. (2007). The sensitivity of yeast mutants to oleic acid implicates the peroxisome and other processes in membrane function. Genetics 175, 77–91.

37. Saleem, R.A., Long-O’Donnell, R., Dilworth, D.J., Armstrong, A.M., Jamakhandi, A.P., Wan, Y., Knijnenburg, T.A., Niemistö, A., Boyle, J., Rachubinski, R.A., et al. (2010). Genome-wide analysis of effectors of peroxisome biogenesis. PLoS One 5, e11953.

38. Aviner, R., Geiger, T., and Elroy-Stein, O. (2014). Genome-wide identification and quantification of protein synthesis in cultured cells and whole tissues by puromycin-associated nascent chain proteomics (PUNCH-P). Nat Protoc 9, 751–760.

39. van Roermund, C.W.T., Waterham, H.R., Ijlst, L., and Wanders, R.J.A. (2003). Fatty acid metabolism in Saccharomyces cerevisiae. Cell Mol Life Sci 60, 1838–1851.

40. Kruppa, M., and Kolodrubetz, D. (2001). Mutations in the yeast Nhp6 protein can differentially affect its in vivo functions. Biochem Biophys Res Commun 280, 1292–1299.

41. Itakura, E., Kishi, C., Inoue, K., and Mizushima, N. (2008). Beclin 1 forms two distinct phosphatidylinositol 3-kinase complexes with mammalian Atg14 and UVRAG. Mol Biol Cell 19, 5360–5372.

42. Hentze, M.W., Castello, A., Schwarzl, T., and Preiss, T. (2018). A brave new world of RNA-binding proteins. Nat Rev Mol Cell Biol 19, 327–341.

43. Hogan, D.J., Riordan, D.P., Gerber, A.P., Herschlag, D., and Brown, P.O. (2008). Diverse RNA-binding proteins interact with functionally related sets of RNAs, suggesting an extensive regulatory system. PLoS Biol 6, e255.

44. Zhang, C., Wang, X., Park, S., Chiang, Y., Xi, W., Laue, T.M., and Denis, C.L. (2014). Only a subset of the PAB1-mRNP proteome is present in mRNA translation complexes. Protein Sci 23, 1036–1049.

45. Crawford, R.A., Ashe, M.P., Hubbard, S.J., and Pavitt, G.D. (2022). Cytosolic aspartate aminotransferase moonlights as a ribosome-binding modulator of Gcn2 activity during oxidative stress. Elife 11.

46. Mayr, C. (2019). 3’ UTRs Regulate Protein Functions by Providing a Nurturing Niche during Protein Synthesis. Cold Spring Harb Symp Quant Biol 84, 95–104.

47. Longtine, M.S., McKenzie, A., Demarini, D.J., Shah, N.G., Wach, A., Brachat, A., Philippsen, P., and Pringle, J.R. (1998). Additional modules for versatile and economical PCR-based gene deletion and modification in Saccharomyces cerevisiae. Yeast 14, 953– 961.

48. Ginestet, C. (2011). ggplot2: Elegant Graphics for Data Analysis. J R Stat Soc Ser A Stat Soc 174, 245–246.

49. Sprouffske, K., and Wagner, A. (2016). Growthcurver: an R package for obtaining interpretable metrics from microbial growth curves. BMC Bioinformatics 17, 172.

50. Wickham, H., Averick, M., Bryan, J., Chang, W., McGowan, L., François, R., Grolemund, G., Hayes, A., Henry, L., Hester, J., et al. (2019). Welcome to the Tidyverse. J Open Source Softw 4, 1686.

51. Lesnik, C., Golani-Armon, A., and Arava, Y. (2015). Localized translation near the mitochondrial outer membrane: An update. RNA Biol 12, 801–809.

52. Tyanova, S., Temu, T., and Cox, J. (2016). The MaxQuant computational platform for mass spectrometry-based shotgun proteomics. Nat Protoc 11, 2301–2319.

53. Zhang, X., Smits, A.H., van Tilburg, G.B., Ovaa, H., Huber, W., and Vermeulen, M. (2018). Proteome-wide identification of ubiquitin interactions using UbIA-MS. Nat Protoc 13, 530–550.

54. Zuba-Surma, E.K., Kucia, M., Abdel-Latif, A., Lillard, J.W., and Ratajczak, M.Z. (2007). The ImageStream System: a key step to a new era in imaging. Folia Histochem Cytobiol 45, 279–290.

55. Cox, J., Neuhauser, N., Michalski, A., Scheltema, R.A., Olsen, J. V, and Mann, M. (2011). Andromeda: a peptide search engine integrated into the MaxQuant environment. J Proteome Res 10, 1794–1805.

56. R Core Team (2014). R: A language and environment for statistical computing. R Foundation for Statistical Computing, Vienna, Austria. *URL* http://www.R-project.org/.

57. Choi, M., Chang, C.-Y., Clough, T., Broudy, D., Killeen, T., MacLean, B., and Vitek, O. (2014). MSstats: an R package for statistical analysis of quantitative mass spectrometry-based proteomic experiments. Bioinformatics 30, 2524–2526.

58. Quast, J.-P., Schuster, D., and Picotti, P. (2022). protti: an R package for comprehensive data analysis of peptide- and protein-centric bottom-up proteomics data. Bioinformatics advances 2, vbab041.

59. Kumar, L., and E Futschik, M. (2007). Mfuzz: a software package for soft clustering of microarray data. Bioinformation 2, 5–7.

60. Yu, G., Wang, L.-G., Han, Y., and He, Q.-Y. (2012). clusterProfiler: an R package for comparing biological themes among gene clusters. OMICS 16, 284–287.

61. Perez-Riverol, Y., Bai, J., Bandla, C., García-Seisdedos, D., Hewapathirana, S., Kamatchinathan, S., Kundu, D.J., Prakash, A., Frericks-Zipper, A., Eisenacher, M., et al. (2022). The PRIDE database resources in 2022: a hub for mass spectrometry-based proteomics evidences. Nucleic Acids Res 50, D543–D552.

